# Genomics of Educational Attainment Across 80 Years of Social and Political Transformation in Germany

**DOI:** 10.64898/2026.09.18.752588

**Authors:** Deniz Fraemke, A. Miller, P. Koellinger, R. Hertwig, D. Richter, S. Zinn, C. Kandler, A. J. Forstner, B. Mönkediek, M. Diewald, A. Teumer, S. E. Baumeister, H. Völzke, U. Völker, H. J. Grabe, L. Bertram, U. Lindenberger, J. Drewelies, S. Kühn, I. Demuth, D. Gerstorf, A. Okbay, P. Biroli, N. Fuchs-Schündeln, K. P. Harden, A. Abdellaoui, M. Malanchini, E. M. Tucker-Drob, Laurel Raffington

## Abstract

Abrupt social and political transformations in twentieth-century Germany may have shifted how polygenic indices relate to educational outcomes across birth cohorts. Following a prespecified analysis plan, we harmonized genomic, educational, and regional data from four German studies, comprising 13,049 individuals born between 1918 and 1995. Polygenic index associations with educational attainment were robust and strengthened modestly across more recent birth cohorts, with no detectable difference between East and West Germany. Associations with intergenerational educational mobility showed no evidence of such strengthening but were stronger in the East than in the West. Polygenic indices were less predictive of educational attainment and educational mobility for women than for men, with no evidence of regional variation. We discuss the methodological limitations of polygenic indices for examining gene– environment interactions, and emphasize that evaluating whether these differences represent progress towards equal educational opportunity requires a normative framework that social science genetics alone cannot supply.

## Introduction

While average levels of education have increased over time, social disparities by geographic region, gender, and parental education in children’s academic skills and educational attainment persist worldwide (e.g., Chmielewski, 2019; Evans et al., 2021; Hu & Qian, 2023). Both genetic and environmental factors are associated with educational trajectories (Abdellaoui et al., 2025; Plomin et al., 1977), yet the interplay between the individual and the culturally specific mechanisms underlying these associations remains poorly understood. Germany offers a unique context in which to examine gene–environment interplay in education, given its historical transformations over the past century, including the division of the country into a state-socialist and authoritarian regime in East Germany and a capitalist democracy in West Germany after the Second World War, and its subsequent reunification in 1990.

Polygenic indices (PGIs) are summary measures of genetic variants previously found to be associated with, for example, educational attainment in over three million people of European genetic ancestry (PGI-Education, Okbay et al., 2022). In one previous finding based on 1,902 individuals from the Socio-Economic Panel genetic subsample, recruited Germany-wide (SOEP-G), the association of this PGI-Education with educational attainment did not differ between East and West Germany before reunification but increased in East Germany thereafter, that is, for those born after 1974 (Fraemke et al., 2025). In East Germany, admission to higher education was politically regulated, with political loyalty, ideological conformity and social class serving as important selection criteria (Below, 2017; Klein et al., 2019). West Germany retained early selection into hierarchically organized school tracks, a structure that tied, and still ties, educational outcomes to family resources (Anweiler et al., 1990; Baier & Lang, 2019). With the reunification of Germany in 1990, the East German educational system was rapidly replaced by the West German institutional framework (Betthäuser, 2019; Klein et al., 2019). Thus, the amplification of PGI-Education associations we observed aligns with theories suggesting that greater social and educational opportunity heightens genetic influences on educational attainment (e.g., Engzell & Tropf, 2019; Heath et al., 1985; Morris et al., 2026). In line with these theories, no regional difference was found before reunification, when both systems constrained access, by political criteria in the East and by family resources in the West. Whether stronger genetic effects are considered desirable depends on the normative model of equality of opportunity that guides social and educational institutions (Grätz & García-Sierra, 2026).

Three key gaps in knowledge remain. First, our earlier study pooled all birth cohorts born before 1975 into a single pre-reunification period, so it is unknown whether PGI-Education associations differed between East and West Germany within that period, particularly in the early years of the division, between 1945 and 1965. During this period, state-socialist East Germany replaced the tripartite school system with ten years of comprehensive schooling and provided preferential access to higher education for students from working-class and farming families (Anweiler et al., 1990; Below, 2017). Second, intergenerational educational mobility refers to differences in educational attainment between parents and their children, indicating whether children attain a higher, lower, or similar level of education than their parents. This relational outcome targets the transmission of education from parents to children, on which the two systems strongly differed (Betthäuser, 2019; Klein et al., 2019). While genomic predictors of education are associated with educational mobility (Belsky et al., 2018), it is unknown to what extent these associations differ between the distinct institutional and educational contexts of former East and West Germany. Third, sociological and economic studies suggest that in contemporary Western industrialized societies, genetic associations with both educational attainment and educational mobility may vary by gender (e.g., Herd et al., 2019; Lahtinen et al., 2023; Rimfeld et al., 2018). East and West Germany differed substantially in gender culture and policy. While the East German government actively promoted women’s economic and educational participation through public childcare and paid study leave, West Germany retained a male-dominated work and educational culture (Below, 2017; Wharton, 1988). Addressing these gaps requires a sufficiently large German genetic dataset with high statistical power, which currently no single German study provides.

This study examines whether and when polygenic index associations with education have remained stable, attenuated, or amplified across the division and reunification of Germany. Following a prespecified analysis plan, we harmonized genomic, educational, and regional data from four German cohorts, comprising 13,049 individuals born between 1918 and 1995 (Figure 1). Our four-cohort mega-analysis probed whether polygenic index associations with educational attainment and intergenerational educational mobility vary by birth year, region (East versus West), and gender. We included three education-genetic indicators: PGI-Education (EA4, Okbay et al., 2022), which can be further split into PGIs for cognitive and non-cognitive skills (Malanchini et al., 2024), to test whether any moderation is specific to one of the two components. PGI-Education, built from the largest discovery sample, is the primary variable of interest. The cognitive and non-cognitive PGIs are secondary measures with smaller expected associations. In addition, we estimated total GREML-based SNP-based heritability of educational attainment in each subgroup.

**Figure 1.**
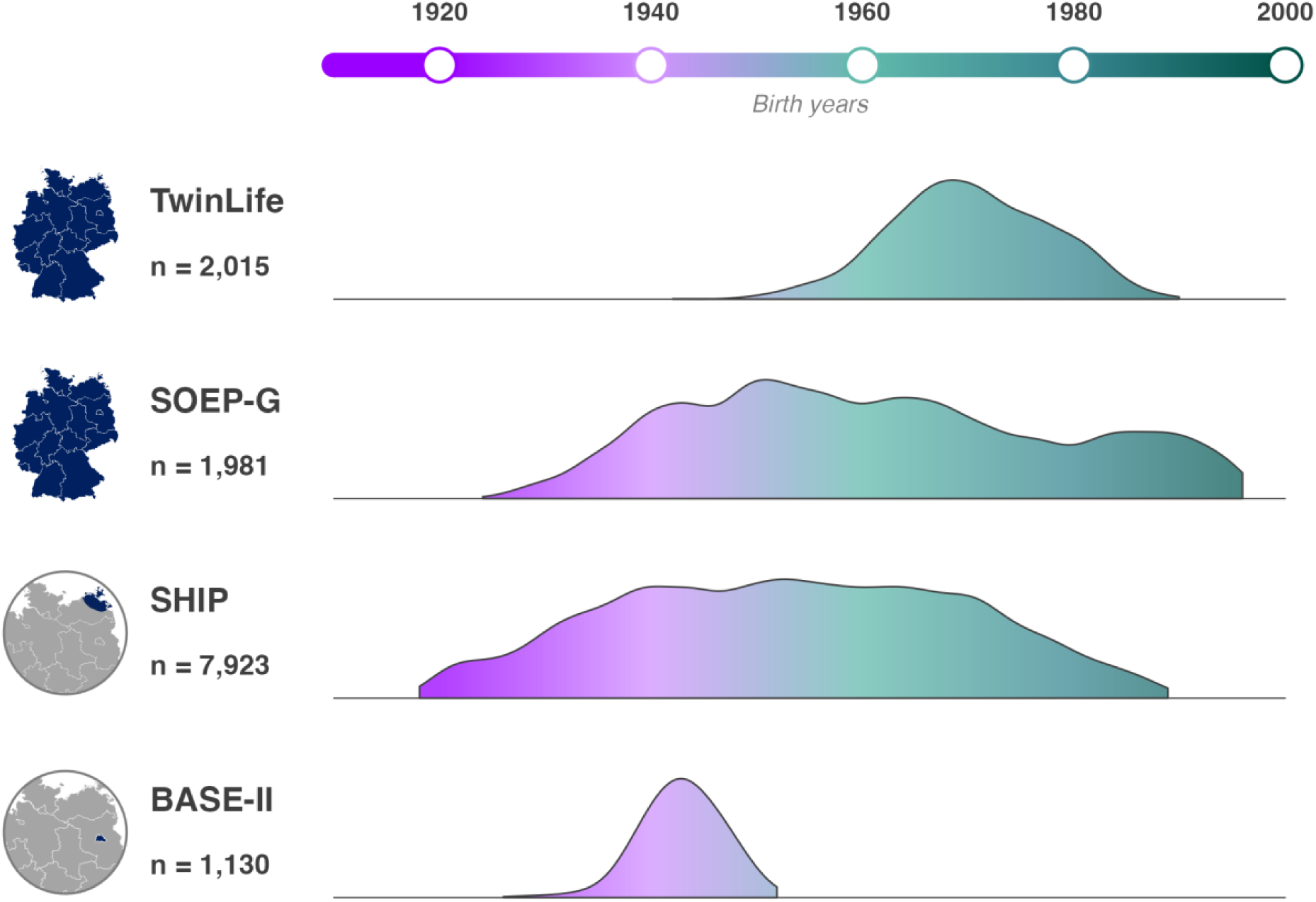
Birth-year distribution and geographic scope of the four contributing studies. The four contributing studies are, from top to bottom, TwinLife, a German twin-family panel study recruited Germany-wide (Mönkediek et al., 2019); SOEP-G, the Socio-Economic Panel genetic subsample, recruited Germany-wide (Goebel et al., 2018; Koellinger et al., 2023); SHIP, the Study of Health in Pomerania, recruited in Mecklenburg-Western Pomerania (Völzke et al., 2022); and BASE-II, the Berlin Aging Study II, recruited in and around Berlin (Bertram et al., 2014). Each ridge shows the distribution of birth years in a study’s educational attainment analytic sample, East and West Germany pooled, as a fixed-bandwidth smooth of the yearly counts; ridge heights are relative frequencies within each study, scaled to the study’s own maximum, and are not comparable across studies. Sample sizes are the corresponding final analytic sample sizes, totalling *n* = 13,049; the intergenerational educational mobility analyses, which additionally required parental education, used *n* = 4,904 participants from TwinLife, SOEP-G, and BASE-II. Only respondents born up to and including 1996 were eligible, which is why the SOEP-G distribution ends abruptly in the mid-1990s. BASE-II was recruited after reunification in a city that had been divided; its older participants were classified as East or West German from their reported residence in the German Democratic Republic, and its younger participants, who were not asked, are not in the analytic sample, which is why the BASE-II distribution ends in the early 1950s. Maps indicate the geographic scope of the contributing studies. The colour gradient along the birth-year axis is a visual scale for birth year and encodes no study characteristic.

## Results

The results are structured into four sections. The first reports descriptive statistics, including regional and gender differences in average education, to help contextualize any differences in the PGI-Education associations across potential moderators. The second section asks whether the association between PGI-Education and educational attainment varies by birth year and region. The third asks the same for intergenerational educational mobility, in the smaller subsample for which parental education was available (*N* = 4,904). The fourth probes whether the PGI-Education associations differed by gender, and whether any gender difference varied over birth year or between regions. Educational attainment is operationalized as relative years of education, that is, years of education standardized over birth year in the pooled sample. Intergenerational educational mobility is the difference between a participant’s educational attainment and the mean of their parents’, each standardized against its own birth-year distribution. Estimates are per 1 *SD* of the predictor. Secondary PGI associations, sensitivity analyses for model dispersion, and leave-one-study-out analyses are reported where applicable.

### 1. Descriptive statistics

The genotyped analytic sample comprised *N* = 13,049 for educational attainment and *N* = 4,904 for intergenerational educational mobility, the latter excluding the Study of Health in Pomerania (SHIP), for which parental education was unavailable (Table 1). East German participants made up *n* = 9,258 of the educational attainment sample, 7,923 of them recruited by SHIP in Mecklenburg-Western Pomerania, so the East German stratum rests largely on one north-eastern state. The overall distribution of educational attainment in the sample was nevertheless broadly in line with figures for the German population, with comparable shares at the top and at the bottom of the educational distribution (Statistisches Bundesamt, 2026a, 2026b).

**Table 1.**
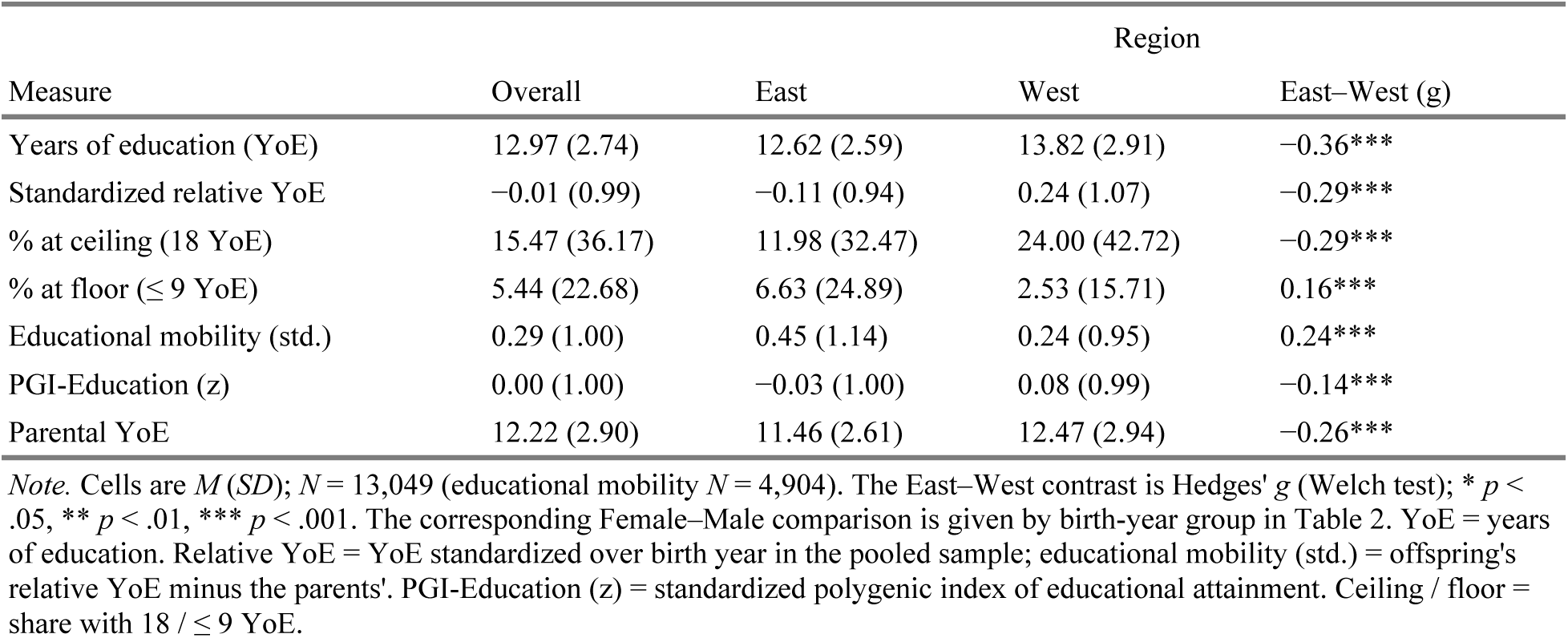
Sample descriptives are reported overall and separately by region.

Mean educational attainment rose by *b* = 0.27 years per decade of birth year, 95% CI [0.24, 0.30], *p* < .001, a trend across the full birth-year range rather than a tabulated contrast. West German participants completed more years of education than East German participants (Hedges’ *g* = −.36, *p* < .001; Table 1), whereas intergenerational educational mobility was higher among East German than among West German participants (*g* = .24, *p* < .001); this regional difference narrowed across birth years (birth year × region *b* = −0.23 per decade, 95% CI [−0.28, −0.18], *p* < .001). Figure 2 shows the shape these differences take. West German participants completed more years of education across most of the birth-year range studied (Figure 2, panel A), while the regional difference in educational mobility reversed, being higher in East Germany among the earliest-born participants and higher in West Germany from approximately the early 1950s onward (Figure 2, panel B). In the subsample for which parental education was available (*N* = 4,904), the association between parental education and educational attainment was stronger in West Germany, *b* = 0.61 (95% CI [0.58, 0.64]), than in East Germany, *b* = 0.39 (95% CI [0.34, 0.45]); the difference was *b* = −0.216, 95% CI [−0.278, −0.154], *p* < .001 (Figure 3, panel B). The strength of this contrast is sensitive to which studies contributed; leave-one-study-out estimates are reported in the Supplementary Results.

**Figure 2.**
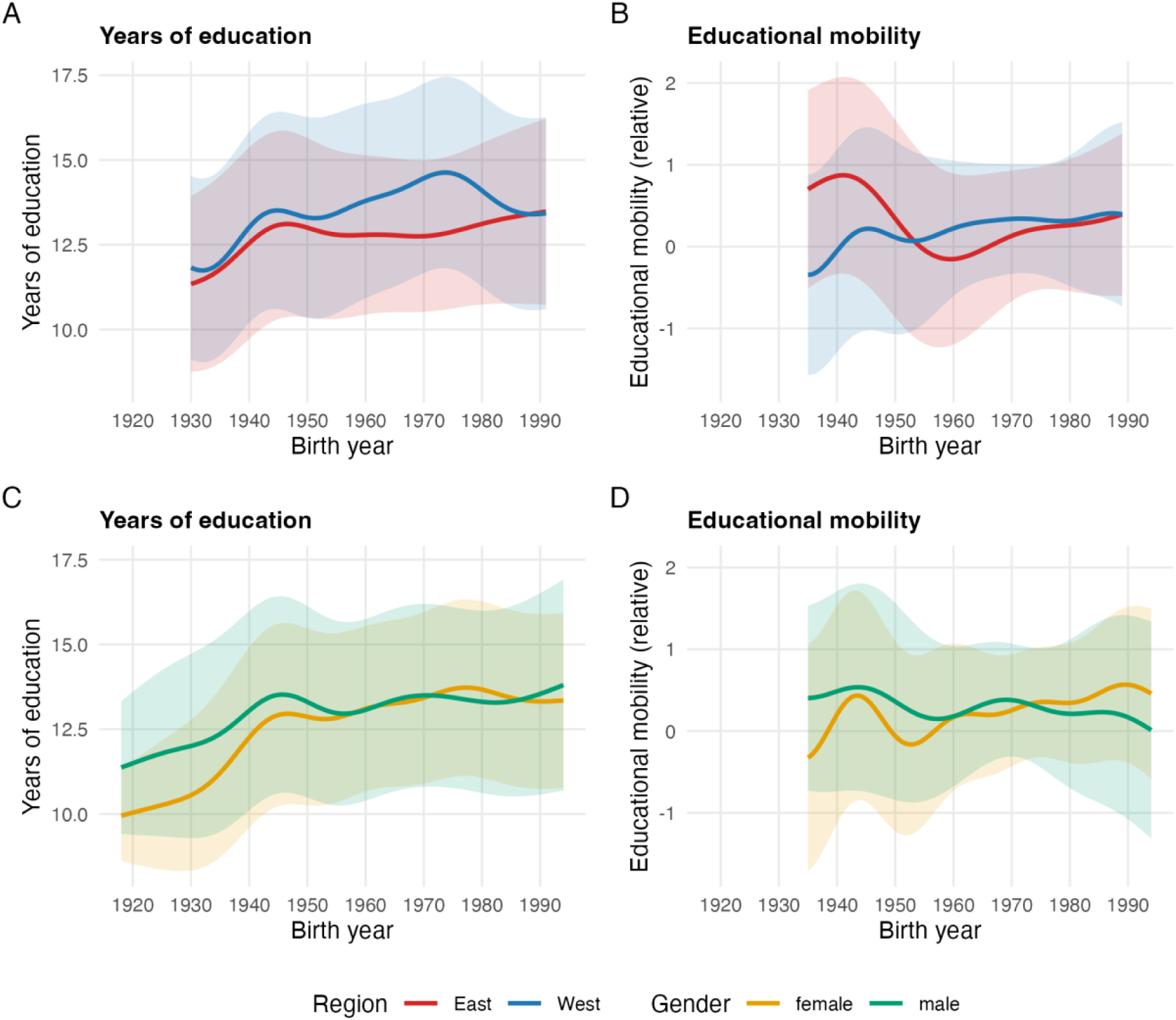
Educational attainment and intergenerational educational mobility across birth year, by region and gender. Panels A and B show educational attainment and intergenerational educational mobility by region, and panels C and D show the same two outcomes by gender. The curves are descriptive local smoothed averages of the outcomes themselves; the PGI-Education associations are reported in the sections that follow. Red curves are East Germany and blue curves are West Germany; amber curves are women and green curves are men. The centre line is the fitted mean and the band around it is that mean plus or minus 1 *SD* of the outcome distribution, which constitutes a dispersion band and not a confidence interval. All four panels share one birth-year axis, and each curve is drawn only over the birth years in which its own group was observed, so a curve stops where that group’s data stop. Figures for educational attainment rely on *n* = 13,049; the educational mobility panels use the smaller sample requiring parental education (i.e., *n* = 4,904).

**Figure 3.**
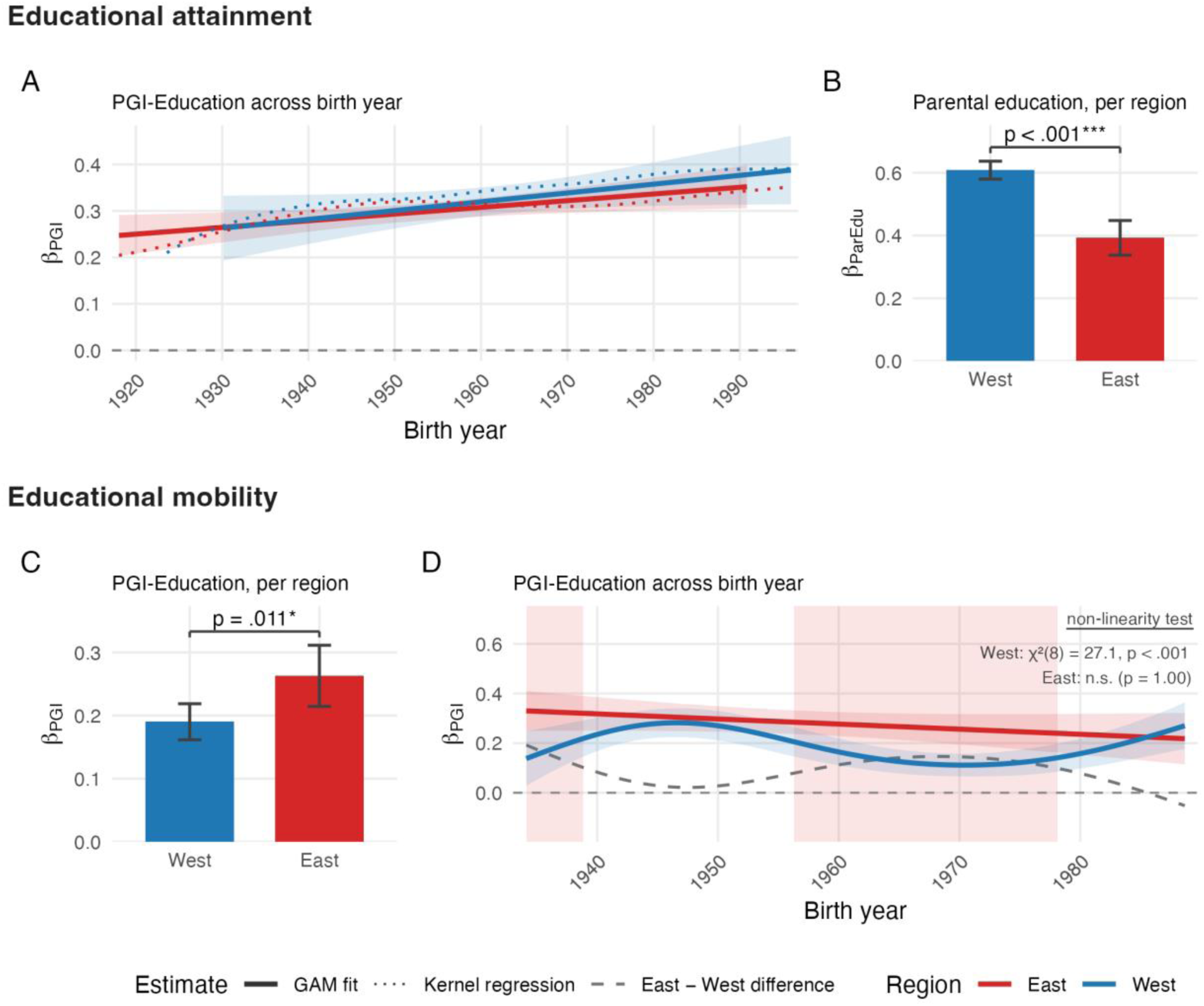
PGI-Education and parental education associations with educational attainment, and PGI-Education associations with intergenerational educational mobility, by region. Panel A shows the PGI-Education slope for educational attainment across birth year in the combined four-study sample. Panel B shows the slope of educational attainment on birth-year-standardized parental education in each region; the East–West contrast it displays is reported in the descriptive statistics. Panel C shows the overall PGI-Education slope for intergenerational educational mobility in each region, and panel D shows that slope across birth year. Red denotes East Germany and blue denotes West Germany throughout. Solid lines are the model-based fits, the dotted lines in panel A are the descriptive kernel-regression overlay, and the dashed grey line in panel D is the East-minus-West difference in the slope, drawn on the same axis as the region curves so that positive values indicate a stronger association in East Germany. Centre values are the estimated slopes, per 1 *SD* of PGI-Education in panels A, C and D and per 1 *SD* of birth-year-standardized parental education in panel B. The bars in panels B and C carry 95% confidence intervals as whiskers, as do the brackets marking the East–West contrasts, and the ribbons in panels A and D are 95% confidence intervals. The shaded vertical windows in panel D mark birth years at which the pointwise 95% confidence interval of the difference excludes zero; the narrower windows from the prespecified band that holds across the whole birth-year range at once, and the per-region non-linearity tests annotated in panel D, are reported in the Results. Panel B reports the association on the educational attainment scale. Because intergenerational educational mobility is a participant’s relative years of education minus the parents’, expressing the same association on the educational mobility scale would subtract exactly 1 from the coefficient, and the East–West contrast is identical under either parameterization. The model does not include interactions of parental education with either birth year or PGI-Education and does not adjust for PGI-Education. Panel A uses the full educational attainment sample, *n* = 13,049. Panels B, C and D use the smaller subsample for which parental education was available, which excludes SHIP, *n* = 4,904; panel B examines educational attainment in that subsample, while panels C and D examine intergenerational educational mobility.

Men had higher average educational attainment and higher intergenerational educational mobility than women (Hedges’ *g* = −.11, *p* < .001, and *g* = −.10, *p* = .002, respectively, a negative *g* indicating the higher male mean). Neither gender difference in means held constant across birth years. The female-minus-male difference in years of education increased by *b* = 0.26 per decade of birth year, 95% CI [0.21, 0.32], *p* < .001, with higher values among men in earlier-born cohorts and among women in more recently born cohorts. A parallel pattern emerged for educational mobility (*b* = 0.12 per decade, 95% CI [0.08, 0.17], *p* < .001; Figure 2, panels C and D). Hedges’ *g* for the Female–Male comparison within three birth-year groups is given in Table 2, and the corresponding East–West contrasts in Supplementary Data S1, Table 2.

**Table 2.** Sample descriptives are reported by gender within each of three birth-year groups, with the Female–Male contrast for each group.

| Measure | Birth year | Gender |  |  |
| --- | --- | --- | --- | --- |
|  |  | Female | Male | Female–Male (g) |
| Years of education (YoE) | < 1950 | 12.08 (2.73) | 12.88 (2.92) | –0.28*** |
|  | 1950–1974 | 13.18 (2.51) | 13.24 (2.71) | 0.00 |
|  | ≥ 1975 | 13.62 (2.60) | 13.37 (2.75) | 0.12* |
| Educational mobility (std.) | < 1950 | 0.22 (1.32) | 0.49 (1.25) | –0.21*** |
|  | 1950–1974 | 0.18 (0.82) | 0.30 (0.82) | –0.15** |
|  | ≥ 1975 | 0.40 (0.80) | 0.18 (1.01) | 0.28*** |
| % at ceiling (18 YoE) | < 1950 | 10.81 (31.06) | 17.24 (37.78) | –0.18*** |
|  | 1950–1974 | 14.83 (35.54) | 17.73 (38.20) | –0.07* |
|  | ≥ 1975 | 17.02 (37.60) | 17.25 (37.80) | 0.00 |
| % at floor (≤ 9 YoE) | < 1950 | 15.17 (35.88) | 6.57 (24.77) | 0.28*** |
|  | 1950–1974 | 1.75 (13.12) | 2.05 (14.16) | –0.03 |
|  | ≥ 1975 | 1.63 (12.65) | 3.09 (17.31) | –0.13* |

*Note.* Female and Male cells are $M(SD)$ within each birth-year group; the third column is Hedges' $g$ (Welch test); \* $p < .05$ , \*\* $p < .01$ , \*\*\* $p < .001$ . A positive $g$ indicates a higher female mean. Educational attainment $N = 5,129$ (< 1950), 6,097 (1950–1974), 1,823 (≥ 1975); the educational mobility rows use the smaller subsample that excludes SHIP. The East–West contrasts for the same groups are reported in Supplementary Data S1, Table 2. PGI-Education and parental education are given overall in Table 1. The 1975 cut marks turning 15 in/after 1990, the year of reunification. YoE = years of education. Relative YoE = YoE standardized over birth year in the pooled sample; educational mobility (std.) = offspring's relative YoE minus the parents'. Ceiling / floor = share with 18 / ≤ 9 YoE.

### 2. The association between PGI-Education and educational attainment strengthened over birth year without evidence of regional moderation

All three polygenic indices were associated with both educational attainment (PGI-Education .31, 95% CI [.30, .33]; PGI-Cognition .18, 95% CI [.17, .20]; PGI-Non-Cognitive .23, 95% CI [.22, .25]) and intergenerational educational mobility (.23, 95% CI [.21, .26]; .13, 95% CI [.10, .15]; .15, 95% CI [.12, .17], respectively), all *p* < .001 (Supplementary Data S3, S6).

The association between PGI-Education and educational attainment was modestly stronger at later birth years (Figure 3, panel A). The pooled PGI-Education × birth year term was *b* = 0.029, 95% CI [0.010, 0.048], *p* = .002, and the same trend modelled as a step at the reunification cutoff was *b* = 0.062, 95% CI [0.014, 0.110], *p* = .011.

Next, we tested whether this cohort increase differed between the former East and West Germany, as previously found in SOEP-G (Fraemke et al., 2025). We found no evidence of such a difference. Neither the PGI-Education × birth year × region interaction, *b* = −0.003, 95% CI [−0.040, 0.035], *p* = .879, nor the PGI-Education × post-reunification × region interaction, *b* = −0.021, 95% CI [−0.116, 0.074], *p* = .667, was significant. The earlier SOEP-G pattern reappeared when the present analyses were restricted to SOEP-G and to that report’s within-SOEP ancestry adjustment, but was absent from the pooled four-study analysis (Supplementary Fig. S1). No regional difference in either the birth-year trend or the change across the reunification cutoff emerged after any single study was omitted (Supplementary Fig. S2; Supplementary Data S2).

We also tested whether the association changed non-linearly across birth year rather than at a constant rate, by letting the PGI-Education slope vary smoothly across birth year. The omnibus comparison provided no support for the nonlinear specification, χ²(16) = 11.94, *p* = .748. Region-specific comparisons likewise found no evidence of departure from linearity in either West Germany, χ²(8) = 0.00, *p* = 1.000, or East Germany, χ²(8) = 12.21, *p* = .142.

We re-estimated the linear model with the prespecified secondary predictors, PGI-Cognition and PGI-Non-Cognitive, in a common analytic sample. The strengthening association with educational attainment over birth year was evident for PGI-Cognition (*b* = 0.023, 95% CI [0.003, 0.042], *p* = .022), but not PGI-Non-Cognitive (*b* = −0.001, 95% CI [−0.020, 0.019], *p* = .955). Neither component showed evidence that this birth-year trend differed between East and West Germany (Supplementary Data S3 and S6).

We then computed the GREML SNP-based heritability of educational attainment, estimating each stratum in its own model from the genetic relatedness among unrelated participants; it was *h*² = .29 (*SE* .04, 95% CI [.22, .36]) across the four region × era strata, East and West Germany before and after reunification. We detected no evidence for heterogeneity across those four strata (Cochran’s *Q* = 0.66, *df* = 3, *p* = .881, *I*² = 0; Supplementary Results and Supplementary Fig. S5).

### 3. The association between PGI-Education and educational mobility was stronger in East Germany than in West Germany

We next asked whether the association between PGI-Education and intergenerational educational mobility differed between regions, in the three studies for which parental education was recorded and therefore excluding SHIP, which contributed only East German participants. The educational mobility analyses exclude SHIP altogether, so unlike the educational attainment analyses this regional comparison is not confounded with contributing study. PGI-Education was more strongly associated with educational mobility in East than in West German participants (PGI-Education × region contrast *b* = 0.073, 95% CI [0.017, 0.129], *p* = .011). The slope was .26 (95% CI [.21, .31]) in East Germany and .19 (95% CI [.16, .22]) in West Germany (Figure 3, panel C).

Three prespecified tests then asked whether this East–West difference was stable across the 1990 reunification divide and over birth year.

For the first test, there was no evidence that the East–West difference in the PGI-Education association with educational mobility changed across reunification. The PGI-Education × post-reunification × region interaction — comparing participants who reached age 15 before 1990 with those who reached age 15 in or after 1990 — was *b* = −0.093, 95% CI [−0.236, 0.051], *p* = .206. The post-reunification group was small and its estimates correspondingly imprecise, so this is a lack of evidence rather than evidence of stability.

For the second test, when the trend over birth year was modelled linearly, there was no evidence that this association varied with birth year overall (*b* = −0.026, 95% CI [−0.055, 0.004], *p* = .085). Nor was there evidence that this linear birth-year trend differed between East and West Germany (*b* = 0.010, 95% CI [−0.048, 0.068], *p* = .732).

For the third test, allowing each region’s trend to bend freely over birth year fit better than modelling it linearly, in a likelihood-ratio comparison of the two specifications, χ²(16) = 27.08, *p* = .041. This curvature was evident only in West Germany, χ²(8) = 27.08, *p* < .001; the East German trend showed no departure from linearity, χ²(8) = 0.00, *p* = 1.000. Modelling the East-minus-West gap directly across birth year, the gap was reliably larger in the East for participants born approximately 1959–1975 (Figure 3, panel D). Across the three tests, then, the East–West difference showed no change across reunification and no linear change over birth year; only the flexible specification located where it was clearest.

Re-estimated with the prespecified secondary predictors in a common analytic sample, the East–West difference in the association with educational mobility was also evident for PGI-Cognition (*b* = 0.082, 95% CI [0.025, 0.138], *p* = .004), but not PGI-Non-Cognitive (*b* = 0.003, 95% CI [−0.053, 0.059], *p* = .928) (Supplementary Data S3 and S6).

Several further checks qualify this contrast. It remained statistically distinguishable from zero in a model that also let the outcome’s spread vary with the predictors (*b* = 0.065, *p* = .035, in the educational mobility subsample), held with the outcome in non-standardized years of education and under a rank transform. It was not distinguishable from zero under an ordered-categorical specification, which carries no family random intercept and collapses the 13-to-18-year step into one category (Supplementary Results). In the leave-one-study-out analyses the contrast remained positive in every reduced sample, attenuated after omitting BASE-II or TwinLife and increased after omitting SOEP, and evidence favouring the nonlinear birth-year specification was attenuated in each reduced sample (Supplementary Fig. S3; Supplementary Data S5 and S6).

### 4. PGI-Education associations with educational attainment and intergenerational educational mobility were weaker among women than men

Our final set of analyses probed gender moderation of polygenic index associations. The association of PGI-Education with educational attainment and intergenerational educational mobility was weaker among women than men (PGI-Education × gender interaction for educational attainment *b* = −0.039, 95% CI [−0.076, −0.001], *p* = .042, and for educational mobility *b* = −0.066, 95% CI [−0.124, −0.007], *p* = .027), and the same ordering held in every region-by-gender cell (Figure 4, panels A and D). Decomposing that interaction, the PGI-Education slope was .29 (95% CI [.27, .32]) among women and .33 (95% CI [.30, .36]) among men.

**Figure 4.**
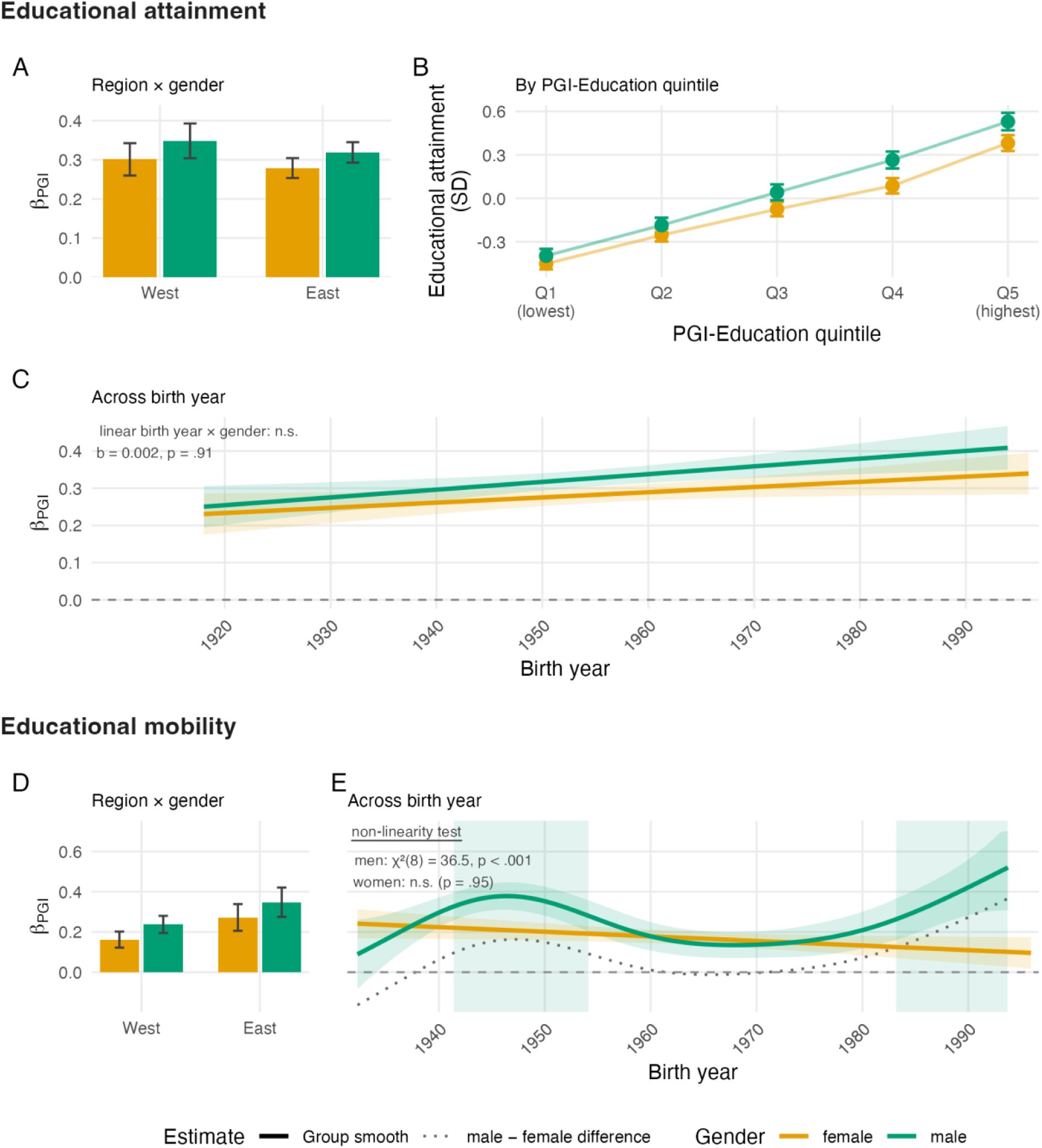
PGI-Education associations with educational attainment and intergenerational educational mobility, by gender. Panel A shows the PGI-Education slope for educational attainment in each region-by-gender cell. Panel B shows mean educational attainment across the PGI-Education distribution by gender, split into PGI-Education quintiles pooled across region and contributing study, quintile 1 being the lowest. It is exploratory and descriptive, and its per-quintile male-minus-female differences are shown in Supplementary Fig. S6. Panel C shows the PGI-Education slope for educational attainment across birth year by gender. Panel D repeats panel A for intergenerational educational mobility, and panel E repeats panel C for that same outcome. Amber denotes women and green denotes men throughout. The gender curves in panels C and E are solid model-based fits, and the dotted grey line in panel E is the male-minus-female difference in the slope, on the same axis as the gender curves. Centre values are the estimated slopes in panels A, C, D and E, per 1 *SD* of PGI-Education, and the raw male and female means of relative years of education in panel B; every error bar, whisker and ribbon is a 95% confidence interval. The shaded vertical windows in panel E mark birth years at which the pointwise 95% confidence interval of the difference excludes zero, drawn in the colour of the gender the difference favours. The narrower windows from the prespecified band that holds across the whole birth-year range at once, and the tests annotated in the panels, are reported in the Results. Sample size for educational attainment is *n* = 13,049 in panels A to C, and *n* = 4,904 for intergenerational educational mobility in panels D and E.

To better understand this interaction, an exploratory follow-up analysis examined whether this gender difference was more pronounced at higher or at lower values of PGI-Education. The male–female difference in mean educational attainment was larger at higher than at lower values of PGI-Education (Figure 4, panel B). Among participants in the top 20% of the PGI-Education distribution, women had completed on average 0.34 years less education than men (*SE* 0.12), against 0.06 years (*SE* 0.09) among those in the bottom 20% (Supplementary Fig. S6). Because the education measure is capped at 18 years, this is likely an attenuated estimate, and the gender difference may be larger at the even higher tail-end of education.

We tested whether this gender interaction reflected a difference in the variance of educational attainment rather than in the association itself. Both interactions remained statistically distinguishable from zero when residual dispersion was modelled, for educational attainment (*b* = −0.044, *p* = .006) and educational mobility (*b* = −0.076, *p* = .002) (Domingue et al., 2021). In the leave-one-study-out analyses both pooled contrasts continued to indicate weaker PGI-Education associations among women in every reduced sample, attenuating most after omitting SOEP (Supplementary Fig. S4; Supplementary Data S2, S3, S5, and S6). Additional exploratory rank and ordinal checks on the gender difference are reported in the Supplementary Results.

We next examined whether this gender difference in polygenic index associations differed across birth years. For educational attainment we detected no such change, whether birth year was modelled linearly (PGI-Education × birth year × gender *b* = 0.002, 95% CI [−0.035, 0.039], *p* = .915) or flexibly (χ²(16) = 19.88, *p* = .226; Figure 4, panel C). For educational mobility the linear interaction was likewise null (*b* = −0.040, 95% CI [−0.099, 0.018], *p* = .178), but a flexible model fit better (χ²(16) = 39.35, *p* < .001), driven by the male trajectory (men χ²(8) = 36.51, *p* < .001; women χ²(8) = 2.78, *p* = .948). Comparing the gender curves directly, the male slope was larger than the female slope among participants born approximately 1943–1951 and 1988–1994, and at no birth year was the female slope larger (Figure 4, panel E). Towards the most recent birth years the male slope rose to its maximum while the female slope fell, so the two curves diverged rather than converged over the period covered.

We detected no regional variation in this gender difference for either outcome. The PGI-Education × region × gender interaction was *b* = 0.017, 95% CI [−0.058, 0.092], *p* = .663, for educational attainment and *b* = 0.013, 95% CI [−0.104, 0.129], *p* = .829, for educational mobility. This is a three-way interaction estimated in region-by-gender cells of unequal size, so it is a lack of evidence for regional moderation rather than evidence that the gender difference was identical in the two regions.

Neither PGI-Cognition nor PGI-Non-Cognitive showed a gender interaction for educational attainment or educational mobility (all *p*s ≥ .294), and at the omnibus level neither component showed the nonlinear gender pattern in educational mobility. The gender difference therefore appeared only for PGI-Education (Supplementary Data S3 and S6).

We detected no evidence for a gender difference in the SNP-based heritability of educational attainment (male .28, *SE* .08; female .38, *SE* .07; Cochran’s *Q* = 0.92, *df* = 1, *p* = .337, *I*² = 0; Supplementary Fig. S5).

As a prespecified negative control, we substituted PGI-Height for PGI-Education to test whether the results were driven by trait-nonspecific properties of a polygenic index. PGI-Height was strongly associated with height, *b* = 0.412, 95% CI [0.399, 0.424], *p* < .001, and weakly but detectably with educational attainment, *b* = 0.025, 95% CI [0.004, 0.047], *p* = .023, and with educational mobility, *b* = 0.040, 95% CI [0.003, 0.078], *p* = .033, uncorrected for the number of sensitivity tests. None of the birth-year, regional or gender moderation patterns observed for PGI-Education and PGI-Cognition emerged for PGI-Height, which is what the negative control was prespecified to test (Supplementary Results; Supplementary Data S3 and S6).

## Discussion

We examined whether polygenic index associations with educational attainment and intergenerational educational mobility varied by birth year, East–West region, and gender in Germany. In sum, we found that these associations were robustly present and strengthened in more recent birth cohorts. While associations with educational attainment showed no detectable regional variation, associations with intergenerational educational mobility were significantly stronger in East than in West Germany. Lastly, we observed a PGI-by-gender interaction, whereby polygenic indices were less predictive of educational attainment for women than for men.

The association between PGI-Education and educational attainment strengthened modestly across successive German birth cohorts. This result broadly replicates trends in the United Kingdom and Sweden (Morris et al., 2026; Pettersson, 2025), while Finnish data show a recent levelling off of this increase (Lahtinen et al., 2023) and Estonian estimates exhibit sensitivity to model specification (Kuznetsov et al., 2026; Rimfeld et al., 2018). This broader pattern of historical amplification may signal a shift towards more meritocratic educational systems, characterized by fewer structural barriers and a greater influence of individual differences. Alternatively, it could reflect the downstream accumulation of social stratification across generations, because population-level polygenic indices pool direct genetic effects, assortative mating, and environmental pathways (Kong et al., 2018; Nivard et al., 2024; Young et al., 2019).

We found no evidence of analogous strengthening over birth years for intergenerational educational mobility. Because educational mobility is measured against a respondent’s own parents, it is less exposed to parental-education stratification than educational attainment (Belsky et al., 2018; Hu & Qian, 2023). Thus, the absence of evidence for amplification across birth cohorts may indicate that the cohort-specific increase in the PGI-Education association with educational attainment reflects intensifying stratification by parental education, which population-level indices partly capture, rather than expanded opportunities for upward educational mobility. In Sweden, the birth cohort trend is also observed within families, where parental environment cannot confound it (Pettersson, 2025). No within-family test was possible here, so which interpretation applies to Germany remains open.

Regarding regional comparisons across former East and West Germany, the association between PGI-Education and educational attainment showed no detectable difference. While a stronger post-reunification association with educational attainment in East Germany is present in SOEP-G data (Fraemke et al., 2025) — and has been statistically replicated both by Spörlein et al. (2025) and our sub-analyses — this pattern did not generalize to our pooled four-study analysis. A similar discrepancy observed in Estonia highlights that context-specific polygenic interactions may be less robust than their main effects (Kuznetsov et al., 2026; Rimfeld et al., 2018). Future studies are needed to resolve whether the results from the SOEP, given its population-representative design, generalize or reflect study-specific nuances.

In contrast, PGI-Education was more strongly associated with intergenerational educational mobility in the former East than in the West. In East Germany, parental education was less predictive of offspring educational attainment, suggesting that dampened stratification by parental education may have allowed the individual differences captured by PGI-Education to play a more prominent role in educational sorting. This aligns with cross-national evidence indicating that while absolute genetic variance in education remains stable across modern societies, its relative share of total variance is higher in more mobile systems (Engzell & Tropf, 2019). While such comparisons have typically been made between different countries, the German division provides a rare within-country test of this hypothesis. The East German state actively suppressed familial privilege through comprehensive schooling and, in its early decades, preferential access to higher education for working-class and farming families, whereas West Germany maintained a more stratified system of early selection (Anweiler et al., 1990; Betthäuser, 2019; Klein et al., 2019; Lahtinen et al., 2024). Birth-year modelling suggested that this East–West difference was clearest among those born between 1959 and 1975, whose schooling coincided with the final decades of the division.

Moreover, our findings indicate a gender disparity in how PGI-Education relates to both educational attainment and intergenerational educational mobility. Whereas women remain less educated than men in most countries (Evans et al., 2021), gender differences in education in Germany are best characterized as a shifting pattern of over- and under-representation across the educational distribution rather than a difference in average years of schooling, which approached parity for cohorts born since 1950 (Hadjar & Berger, 2010; Hu & Qian, 2023). At the lower educational tail, women of recent birth cohorts are under-represented, accounting for roughly 41% of individuals leaving school with only a basic qualification. In the upper-middle range, women show a slight over-representation, earning 53% of Bachelor’s and 51% of Master’s degrees (Statistisches Bundesamt, 2025b). A stark “leaky pipeline” remains at the highest academic tiers, where women earn about 46% of doctorates, up from under 30% in 1992, but hold only 24–30% of German professorships (Gemeinsame Wissenschaftskonferenz, 2012, 2024; Statistisches Bundesamt, 2025a, 2025b).

Our polygenic analyses extend this picture. Even when accounting for baseline gender differences in educational attainment, “PGI-matched” women attain fewer years of education and exhibit lower upward educational mobility than their male counterparts, a difference concentrated among those with high PGI-Education values. We found no evidence that this PGI-by-gender interaction on educational attainment varied significantly by region or across birth years, even as women’s average educational attainment came to exceed men’s, though we caution that this absence of evidence may be an artefact of limited statistical power. Nevertheless, this lack of regional variation suggests that East Germany’s historic institutional efforts to promote women’s educational participation may have been insufficient to dismantle deep-seated gendered structures (e.g., Below, 2017; Hadjar & Berger, 2010; Wharton, 1988). Beyond Germany, similar gender disparities in polygenic index associations have been documented in the United States, Finland, and Estonia, though not in Britain (Herd et al., 2019; Lahtinen et al., 2023; Morris et al., 2026; Rimfeld et al., 2018). Taken together, these patterns support the interpretation that widespread gendered environmental constraints, such as social norms and role expectations, unequal childcare responsibilities, implicit bias, and patriarchal academic cultures, disproportionately divert women with high PGI-Education values away from advanced educational tracks compared to men with the same values.

Multiple conceptions of inequality of opportunity consider gender moderation of genetic effects on education to be reflective of injustice (Grätz & García-Sierra, 2026; Harden, 2021). Theories diverge, however, on what pattern of genetic associations would reflect a just scenario. A fair equality of opportunity framework would hold that just social arrangements are those in which individuals with the same talent and willingness to use it have identical prospects of success, regardless of their social position (Miller, 1999; Rawls, 1971). Alternatively, a functionalist framework posits that individuals possessing specific talents should be matched to corresponding educational and occupational pathways not necessarily to maximize benefit to those individuals but that of society at large (e.g., Parsons, 1991). Both perspectives would welcome the dismantling of gendered barriers so that women’s genetic expression matches men’s, the former to achieve individual equality and the latter to optimize macro-structural utility. Proponents of luck egalitarianism, in contrast, treat any factor outside an individual’s conscious control, including a genetic disposition towards higher education, as a fundamental source of unjust inequality (e.g., Cohen, 2008). Under such a framework, a just society is one that not only minimizes differences in genetic effects across gender but minimizes genetic effects on life outcomes in absolute terms. For example, a luck egalitarian framework would treat the genetic correlation between low PGI-Education scores and ADHD as a signal that schools may be failing to meet the educational needs of neurodiverse children (Cheesman et al., 2025; Demontis et al., 2023). Societal progress therefore depends on a normative framework that genetic data alone cannot supply.

We discuss three limitations of our study. First, polygenic indices are summary measures of genetic correlates normed to a discovery genome-wide association study. Thus weaker associations across different social and political systems, in earlier birth cohorts, or in one gender could reflect social, cultural, demographic, or phenotypic mismatch between our target sample and the discovery sample. A related issue is that PGI-Education carries a substantial environmental signal and is substantially attenuated in within-family estimates (Okbay et al., 2022; Tan et al., 2024). Consequently, polygenic estimates include direct genetic effects and confounding factors. The birth-year amplification of PGI-Education associations may be driven by increasing social stratification of the population over time, also known as social dynastic effects (Nivard et al., 2024). Such stratification is less likely to account for the observed PGI-by-gender interactions, given that gender can be treated as an exogenously and randomly allocated factor within families, which is expected to mitigate stratification bias. Larger, family-based studies with broader geographic coverage are required to determine whether these trends are driven by institutional barriers or by these confounding pathways, and whether either varies across birth cohorts and regions.

Second, testing higher-order interactions with polygenic indices demands considerable statistical power; therefore, the null results reported here should be interpreted as a lack of evidence rather than evidence of absence. Current polygenic indices capture only a fraction of total heritability and are predominantly trained on individuals of European genetic ancestry from Western societies, so they may lack the sensitivity to detect subtle gene–environment interactions, although their main associations, estimated across a vast range of individuals, contexts, and birth cohorts, remain informative. Methods that account for measurement error in polygenic estimates are on the horizon (e.g., Frach et al., 2026), but it remains unclear whether they will increase sensitivity to environmental contexts or primarily sharpen main associations that are stable across recent Western societies.

Third, our sample composition and our measurement of region introduce potentially important imprecision. The four contributing studies differ in setting and coverage, and two of them were geographically localized with restricted birth-year ranges, so study-specific differences may mask or generate apparent birth-year and regional dynamics, and the East German stratum of the educational attainment sample rests largely on one north-eastern state. Germany’s division and reunification do not constitute a clean natural experiment (Becker et al., 2020). Region was not randomly assigned, and pre-separation differences are documented along some dimensions. Migration patterns, differential survival, which favours the healthier and more educated among the oldest birth cohorts assessed late in life, and recruitment biases in the contributing studies actively shape who is observed in each region over time.

In conclusion, our investigation demonstrates that polygenic index associations with educational attainment and intergenerational educational mobility are both robustly present and appear to be shaped by birth cohort, region, and gender. Across multiple European countries, polygenic indices are increasingly predictive of educational outcomes over successive birth cohorts, yet they remain less predictive for women than for men. Judging whether societal progress will best be represented by gender-equalized, strong genetic associations — or, conversely, by weaker genetic effects overall that diminish the impact of the genetic lottery — rests on a normative framework that the field of social science genetics alone cannot supply.

## Methods

We combined genomic, educational, and regional data from four German studies in an individual-level mega-analysis (cf. Tropf et al., 2017). DNA processing, polygenic-index construction, and phenotype coding were harmonized before model specifications were applied to the pooled data. The design, hypotheses, estimators, and inferential criteria were set out in a public analysis plan. Prespecified, exploratory, and additional analyses are distinguished throughout; departures from the plan and their rationale are catalogued in the deviations record in the public analysis repository.

### Study Design and Participants

#### Participating studies

The Socio-Economic Panel genetic subsample (SOEP-G) is a genotyped subsample of the population-based, multigenerational Socio-Economic Panel (Goebel et al., 2018; Koellinger et al., 2023). TwinLife is a representative, cross-sequential German twin-family panel (Mönkediek et al., 2019). Only its adult and parental generation contributed to the present analyses. The Berlin Aging Study II (BASE-II) is a multidisciplinary Berlin study comprising older and younger adult samples (Bertram et al., 2014). The Study of Health in Pomerania (SHIP) comprises the population-based SHIP-START and SHIP-TREND study samples recruited in Mecklenburg-Western Pomerania (Völzke et al., 2022). Per-study birth-year distributions are shown in Figure 1, and descriptive statistics by study in Supplementary Data S1.

Genotyping used Illumina Global Screening Arrays in SOEP-G and TwinLife, Affymetrix Genome-Wide Human SNP Array 6.0 in BASE-II and SHIP-START, and Illumina HumanOmni2.5 and Global Screening Array batches in SHIP-TREND. Each study supplied genotypes that its own data provider had already imputed on genome build GRCh37, against the Haplotype Reference Consortium panel in BASE-II and version 1.1 of that panel in SHIP-START, SHIP-TREND, and SOEP-G, and against 1000 Genomes Phase 3 (v5) in TwinLife. Pre-imputation quality control was performed by the providers and differed across studies. Comparability across the five genotype datasets therefore rests on the harmonization applied here, which is described under Genotype harmonization and polygenic scoring below.

#### Eligibility and analytic samples

We included respondents born up to and including 1996. This fixed birth-year ceiling replaced the planned study-specific age-at-observation criterion and excluded respondents who might not yet have completed their education (OECD, 2024). Analyses were restricted to respondents classified as genetically similar to European reference populations, matching the ancestry composition of the discovery genome-wide association studies (GWAS) (Price et al., 2006). Respondents were projected onto principal components estimated in the 1000 Genomes phase 3 reference panel, and were retained where the nearest of the non-admixed reference centroids — European, African, East Asian, and South Asian — was the European one, and the Mahalanobis distance to that centroid fell within a chi-square cutoff of .999 on six components. The classification was applied within each genotype dataset, before the studies were merged. SOEP-G is the exception. It carried over a European-ancestry keep-list computed for that collection in earlier work; that list’s projection is not documented, and the SOEP-G genotypes are no longer available to re-project. Because the list was built on a more stringently quality-controlled release than the one scored here, respondents absent from the earlier release were retained alongside the listed Europeans, and only listed respondents classified as non-European were excluded. Respondents also required non-missing birth year, educational attainment, region, recorded gender, and PGI-Education. Observations with a pooled-sample standardized PGI-Education outside ±3 *SD* were excluded as an outlier safeguard.

TwinLife twins and non-twin siblings were excluded so that TwinLife contributed only its adult and parental generation, which covered the historical birth-year range of interest. Only a small number of twins and non-twin siblings were aged 25 or older and otherwise eligible. Multiple parents, spouses, or other adult relatives from the same TwinLife families remained eligible and were kept in the analytic sample rather than reduced to one adult per family. SOEP-G respondents were not screened for shared household membership; as in TwinLife, adults from the same household would have remained eligible. SHIP contributed to the East-German stratum only, with SHIP-START respondents identified as West residents in 1989 excluded (*n* = 57) and SHIP-TREND respondents assigned East because no corresponding pre-1989 location measure was available.

The educational attainment analytic sample comprised *N* = 13,049 respondents with observed years of education and PGI-Education. The analytic sample for intergenerational educational mobility additionally required parental education and comprised *N* = 4,904. Parental education was unavailable in SHIP, which therefore did not contribute to educational mobility analyses. Models used complete observations for all variables required by the respective specification; no outcome or covariate values were imputed. Contributing-study and analytic-sample composition are reported in the descriptive tables (Supplementary Data S1).

### Measures

#### Polygenic indices

The focal polygenic index was PGI-Education, based on the EA4 GWAS of educational attainment in approximately three million individuals of European genetic ancestries (Okbay et al., 2022). Posterior SNP weights were estimated with SBayesR (Lloyd-Jones et al., 2019) and applied with allele-aware PLINK2 scoring (Chang et al., 2015). Every polygenic index in every contributing study was built by the same procedure, described below.

To limit sample-overlap bias, all contributing studies were scored from EA4 summary statistics excluding SHIP, as using one weight set for every study also kept the studies on a common scoring scale. BASE-II, which contributed approximately 0.07% of the EA4 discovery sample, was scored from the same weights, because the expected overlap inflation at that share is negligible. The departure from the planned study-specific leave-one-out scoring is documented in the deviations record.

The prespecified secondary polygenic indices were PGI-Cognition and PGI-Non-Cognitive, using published cognition and non-cognitive weights previously applied in this combination (Demange et al., 2021; Lee et al., 2018; Malanchini et al., 2024). The published PGI-Non-Cognitive weights were used in all contributing studies. PGI-Height was included as a negative-control predictor (Yengo et al., 2022).

#### Genotype harmonization and polygenic scoring

The five genotype datasets were harmonized before scoring. Variant identifiers were normalized to chromosome and position, and where that left duplicate records, including split multiallelic sites, only the first was retained. Within each dataset, variants were then filtered on imputation quality at *R*² ≥ .3; in SHIP-TREND, whose two genotyping batches were imputed separately, a variant had to clear the threshold in both. The datasets were then restricted to the intersection of their remaining variants, 6,878,689 SNPs carrying identical identifiers in every dataset. Because TwinLife was imputed against the smaller 1000 Genomes panel, its variant catalogue determined that intersection; restricting every study to it managed the panel difference rather than removing it, but it ensured that no study contributed index variants the others lacked.

Posterior SNP weights were estimated with SBayesR as implemented in GCTB 2.5.2, using a four-component mixture with variance-class scaling factors of 0, .01, .1, and 1 and mixing proportions of .95, .02, .02, and .01, and a chain of 10,000 iterations of which 2,000 were discarded as burn-in; the linkage-disequilibrium reference was the set of shrunk sparse matrices estimated in 50,000 European-ancestry UK Biobank participants, and the major histocompatibility region was excluded. Indices were computed in PLINK 2.0 from hard-called genotypes, with the effect allele of each weight matched to the genotype internally. Twenty ancestry principal components were estimated on the merged genotypes of all five datasets, restricted to autosomal variants with minor allele frequency ≥ .05, missingness ≤ .02, and Hardy-Weinberg equilibrium *p* ≥ 1 × 10⁻⁶, with long-range linkage-disequilibrium regions excluded and the remaining variants pruned to approximate independence (*r*² < .05 in sliding 1,000-variant windows). So that family structure did not enter the leading components, they were estimated in an unrelated subset of the pooled sample (KING-robust kinship < .0884, approximately third-degree relatives) and then projected onto all participants.

Within each relevant pooled analytic sample, each polygenic index was residualized on the first 10 of these components before analysis (Abdellaoui et al., 2013; Price et al., 2006). The residualized scores were then standardized within that analytic sample, so model coefficients represent a one-*SD* difference on a common pooled-sample scale.

#### Educational attainment

Educational qualifications recorded in the four contributing studies were harmonized onto a common full-time years-of-education scale. This harmonization aligned study-specific qualification measures; educational attainment was subsequently standardized over birth year in the pooled analytic sample. The common scale ranged from 7 years for a compulsory-school qualification to 18 years for a university degree; tertiary duration was added to secondary schooling, including 1.5 years for vocational training and 5 years for a university degree.

The focal educational attainment outcome expressed each respondent’s years of education relative to people born in nearby years. For each birth year, we estimated a Gaussian-kernel-weighted local mean and standard deviation over birth year (bandwidth = 5 years) and standardized educational attainment against these local means and standard deviations. The kernel was estimated in the pooled sample and borrowed more information from adjacent than distant birth years, avoiding discontinuities from hard birth-year bins. Kernel-standardized scores required a local effective sample size of at least 10 and an estimable local standard deviation. Because the outcome is standardized within birth year, a change across birth years in the dispersion of years of education would itself change the scale on which the association is measured; sensitivity analyses varying the outcome scale and the estimator are described below.

#### Intergenerational educational mobility

Intergenerational educational mobility was the difference between the respondent’s educational attainment and their parents’, each standardized against its own birth-year distribution (Belsky et al., 2018):

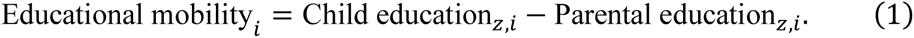

Parental education was calculated as the mean of all available maternal and paternal years of education. When education was available for only one parent, that parent’s value was used; respondents without education information for either parent were excluded from the educational mobility analyses. Parental birth year was calculated the same way, as the mean of the maternal and paternal birth years the harmonized data held, and was approximated as the respondent’s birth year minus 28 years where neither was available. Mean parental education was then standardized with the same five-year Gaussian kernel over parental birth year. Positive educational mobility values indicate that the respondent’s birth-year-relative educational position exceeds that of the parents; negative values indicate the reverse.

#### Birth year, region, and recorded gender

Birth year was used continuously, retaining decimal values (e.g., 1989.43) rather than rounding to completed calendar years. It was standardized in the linear interaction models and mean-centred, but left in year units, in the generalized additive models (GAMs). A reunification indicator distinguished respondents who reached age 15 before 1990 from those who reached age 15 in or after 1990; equivalently, respondents born in 1975 or later were assigned to the post-reunification group. The age-15 cutoff classified respondents according to whether their formative school years extended into the post-reunification period and followed the age-at-transition approach of Rimfeld et al. (2018).

Region represented formative attachment to East or West Germany and was derived separately within each contributing study from the available historical information. In SOEP-G, assignment required a consistent East or West residence through the respondent’s schooling window, reconstructed from 1989 residence and annual household region. In BASE-II, the older sample was classified from a self-report of having lived at least one year in the German Democratic Republic (GDR; East Germany), which records neither when that residence fell nor how long it lasted; unambiguous reported GDR or Federal Republic of Germany (FRG; West Germany) schooling was used when that item was missing. The item was asked of the older sample only, so the younger BASE-II sample could not be classified, and because an observed region was required of every respondent, it did not enter any analytic sample. In TwinLife, pre-1990 births were classified primarily from reported GDR birth and otherwise from first observed federal-state residence; post-1990 births were classified from first observed residence. SHIP’s treatment is described under eligibility above. Mixed or indeterminate regional histories were set to missing for the focal East–West comparison.

Region and recorded gender were effect-coded as −0.5 for West and +0.5 for East, and −0.5 for male and +0.5 for female. This coding centres lower-order terms at the unweighted midpoint of the groups while leaving the focal higher-order interactions and model predictions unchanged. Gender was self-reported by respondents in the contributing studies’ questionnaires, which supplied only binary gender/sex categories and therefore did not capture non-binary gender identities.

### Statistical Analyses

Analyses were conducted in R, with focal linear and GAM specifications fitted using mgcv::bam (Wood, 2017, 2025). Tests were two-sided with α = .05, and estimates are reported with 95% confidence intervals.

Eligible adults from the same TwinLife family were retained, and their dependence was accounted for by adding a family-level random intercept to each focal model (s(fid_re, bs = “re”)). A sensitivity analysis repeated the focal models after retaining one adult per TwinLife family; the estimates are reported in Supplementary Data S3 and S6.

Contributing study was not adjusted for in the models of polygenic index associations. SHIP contributed to the East-German stratum only, and the studies differed in the birth years they covered, so study indicators and study-by-index terms were collinear with the regional and birth-year contrasts these analyses estimated; adjusting for study would remove part of the variation of interest. The SNP-based heritability models described below did include contributing-study fixed effects, because there the estimand was a variance decomposition and between-study differences in mean educational attainment would otherwise enter the variance components.

#### Descriptive PGI associations

We first described the adjusted main association of PGI-Education, PGI-Cognition, and PGI-Non-Cognitive with both educational outcomes. For educational attainment, ordinary least-squares models adjusted for birth year, region, and recorded gender; the educational mobility models additionally adjusted for parental birth-year-standardized education. These associations were descriptive, not tests of historical moderation.

#### Birth-year and regional variation in PGI associations

The prespecified analyses examined whether the association between PGI-Education and each educational outcome varied jointly by birth year and region. The linear specification included PGI-Education, birth year, region, recorded gender, and all interactions among these variables through the three-way level. The focal term was PGI-Education × birth year × region. A parallel step-function model replaced birth year with the reunification indicator, making PGI-Education × reunification × region the focal term.

For educational mobility, these models additionally controlled parental birth-year-standardized education with the interaction structure specified in the analysis plan: its main effect, its two-way interactions with PGI-Education, birth year or reunification, and region, and the corresponding three-way terms. This allowed the parental-education association to vary across the same historical and regional dimensions as the focal polygenic index association.

We evaluated nonlinear birth-year variation with varying-coefficient GAMs (Wood, 2017, 2025). For each educational outcome, three nested specifications represented (a) region-specific PGI associations that were constant across birth year, (b) region-specific linear changes in the PGI association across birth year, and (c) the same linear terms plus region-specific smooth changes across birth year. The nonlinear specification used thin-plate regression splines with basis dimension *k* = 8, fitted by fast restricted maximum likelihood; cubic regression splines were a basis-function sensitivity check. Basis adequacy was evaluated with the *k*-index. Because smoothness was penalized and selected from the data rather than fixed in advance, each region-specific smooth was shrunk toward its linear term unless the data supported curvature; that penalization, together with the nested comparisons and the basis-function check, was what distinguished supported curvature from overfitting.

The linear PGI-Education × birth year × region interaction was evaluated in the linear model. Nonlinearity was evaluated by a nested likelihood-ratio comparison of the linear and nonlinear GAMs, with AIC and BIC used as supporting fit indices. Approximate smooth *F* tests were retained as diagnostics but did not determine whether a trajectory was nonlinear, because they combine linear trend and curvature. Cell-specific departures from linearity were evaluated with clean nested comparisons that added only the focal region’s smooth to the linear model. Simultaneous 95% confidence bands, which provide joint coverage across the full birth-year range rather than separate coverage at each birth year, were used to identify intervals in which the East–West difference in the PGI association differed from zero. The figures show pointwise 95% confidence intervals at each birth year; inference about intervals of East–West difference was based on the simultaneous bands.

The educational mobility GAMs controlled parental education at the same functional flexibility as PGI-Education. Alongside its main and regional effects, separate parental-education smooths over birth year were estimated for East and West.

The regional comparison of the association between parental education and educational attainment was estimated separately, as educational attainment on parental education × region, adjusted for birth year and recorded gender with a per-family random intercept.

#### Gender variation in PGI associations

Prespecified linear models for both educational outcomes added the four-way PGI-Education × birth year × region × recorded gender interaction and all constituent lower-order terms. The GAM extension estimated PGI-Education slopes over birth year separately by recorded gender with regions pooled and then separately in each of the four region × recorded-gender cells. Educational mobility models again controlled parental education at the same cell-specific functional flexibility as PGI-Education. The planned logistic-transition model did not converge and is not reported.

#### Robustness, sensitivity, and exploratory analyses

The focal region and region × recorded-gender GAMs were refitted after omitting each contributing study in turn. The composition of each reduced sample is given in Supplementary Table S1. Two further sensitivity analyses were conducted unconditionally. First, the prespecified secondary-predictor analyses repeated the focal linear and four-way gender models with PGI-Cognition and PGI-Non-Cognitive. The Benjamini-Hochberg procedure controlled the three-score family of statistical tests comprising PGI-Education, PGI-Cognition, and PGI-Non-Cognitive at *q* = .05 (Benjamini & Hochberg, 1995). That family covers the substitution by the secondary predictors. Contrasts falling outside it, including the secondary-predictor East–West educational mobility contrast, were evaluated on uncorrected *p* at *α* = .05, as were the prespecified gender contrasts, which were specified at that level in the analysis plan rather than as part of the corrected family. The planned within-family PGI-Education was not computed, because the corresponding summary statistics were unavailable (Tan et al., 2024). Second, PGI-Height served as a negative-control predictor in models of educational attainment and educational mobility. Its association with measured or reported height was examined separately as a positive-control check that the score captured its intended phenotype (Lipsitch et al., 2010). An exploratory analysis tested whether parental education moderated the PGI-Education association and whether that moderation differed by region (Supplementary Fig. S7).

Following the analysis plan, heteroscedasticity analyses were conditional on evidence from a focal interaction test. Heteroscedasticity was examined with Gaussian location-scale models that allowed residual dispersion to vary with birth year, the focal PGI, or their interaction (Domingue et al., 2021). For each focal interaction that was significant in the primary random-effects analysis, we re-estimated the corresponding mean model under constant residual dispersion and under a dispersion model containing the relevant focal predictors and their interactions. A likelihood-ratio test assessed whether modelling heteroscedasticity improved model fit, and robustness was evaluated by comparing the focal interaction estimate and its *p* value between the constant- and varying-dispersion models. Because this location-scale specification could not include the family random intercept, it used the full-family sample without that term and the focal random-effects result only as the decision gate.

Additional analyses that were not part of the prespecified plan varied the outcome scale, the estimator, the scope of the ancestry residualization applied to PGI-Education, and the ancestry adjustment of the outcome model itself, which added ancestry principal components as main effects and as interactions with the moderators (Supplementary Tables S2–S4).

#### SNP-based heritability

We estimated SNP-based heritability of educational attainment in observed years of education with genome-based restricted maximum likelihood (GREML) in the GCTA software (Yang et al., 2011). The prespecified strata crossed region with the pre-/post-reunification indicator, and each stratum was estimated in its own model rather than by splitting a single pooled fit; because the strata contained disjoint respondents, no respondent contributed to more than one estimate. The primary estimator used study-specific genetic-relatedness matrices, constructed from autosomal variants with minor allele frequency ≥ .01 and a within-study relatedness cutoff of .05, and combined them block-diagonally. Contributing-study indicators were included as fixed effects so that between-study differences in mean educational attainment did not enter the variance components. Models additionally adjusted for recorded gender, 20 ancestry principal components calculated across contributing studies, and linear and quadratic birth-year terms.

Because the strata contained disjoint respondents, equality of SNP-based heritability across the four region × reunification cells was evaluated with Cochran’s *Q* applied to the cell-specific estimates rather than the planned joint multi-GRM likelihood-ratio test. Pairwise *z* tests and *I*² described the location and magnitude of heterogeneity (Cochran, 1954; Higgins et al., 2003). A pooled cross-study genetic-relatedness matrix with a .20 cutoff and separate study-specific estimates served as sensitivity analyses. Region-only, recorded-gender-only, and region × recorded-gender strata were exploratory. Design-based power was calculated from the empirical variance of off-diagonal genetic relatedness following Visscher et al. (2014).

### Use of Generative Artificial Intelligence

During the preparation of this work the authors used OpenAI’s ChatGPT and Anthropic’s Claude to assist with language editing and with analysis and figure code. The authors reviewed and edited all output and take full responsibility for the content of this article.

## Declarations

### Analysis Plan

The design, hypotheses, estimators, and inferential criteria were specified in a publicly available analysis plan (https://osf.io/h4sg8/). Because some contributing data had been analysed previously, the plan is not a preregistration.

### Data Availability

Data availability differs across the four contributing studies, and each statement below is that of the study that owns the data.

**Study of Health in Pomerania (SHIP).** The data of the SHIP study cannot be made publicly available due to the informed consent of the study participants, but it can be accessed through a data application form available at https://fvcm.med.uni-greifswald.de/ for researchers who meet the criteria for access to confidential data.

**TwinLife.** TwinLife survey data are curated and distributed via the GESIS Data Catalogue (https://search.gesis.org/research_data/ZA6701) and are available for academic research and teaching upon completion of the GESIS/TwinLife Data Use Agreement. TwinLife genetic data is available to researchers upon completion of an additional Data Use Agreement with the TwinLife Research Data Centre at Bielefeld University. Comprehensive study documentation, codebooks, and a short guide are publicly available (https://www.twin-life.de/documentation). For enquiries, please contact.

**Socio-Economic Panel genetic subsample (SOEP-G).** All data used in this study are fully accessible to eligible researchers upon application to the management of DIW Berlin (https://www.diw.de/en/diw_01.c.601584.en/data_access.html, contact). All questionnaires and summary statistics of each item are provided in the data documentation of the SOEP Innovation Sample (https://paneldata.org/soep-is/). DIW Berlin shares the genetic principal components and all polygenic indices constructed for SOEP-G in a standard phenotype file, and handles data access applications for the raw genetic data.

**Berlin Aging Study II (BASE-II).** The data presented in this study are available from the BASE-II office, please see https://www.base2.mpg.de/7549/data-documentation for details. Towards that end, we have established procedures over the past ten and more years that we have successfully implemented literally hundreds of times. We are not in a position to make data publicly available because these contain information that could compromise research participants’ privacy and consent.

### Code Availability

All analysis code is version-controlled in a repository (https://github.com/denizFraemke/GxE-education-germany) that is made public on publication of the preprint. The estimates reported here were invariant across the code revisions made during the analysis period.

### Ethics Approval

Ethics approval was obtained separately by each contributing study.

**Study of Health in Pomerania (SHIP).** The study was allowed under the recommendations of the Declaration of Helsinki. The medical ethics committee of the University of Greifswald approved the study protocol, and oral and written informed consents were obtained from each of the study participants.

**TwinLife.** The TwinLife study was reviewed and approved by the German Psychological Society (Deutsche Gesellschaft für Psychologie; protocol number RR 11.2009). The Ethics Committee of the Medical Faculty of the University of Bonn (No. 113/18) reviewed and approved the protocols for the molecular genetic analyses via saliva samples. Informed written consent (for molecular genetic data) and informed verbal consent (for phenotypic data) were obtained from all participants and the participant’s legal guardian (for minors).

**Socio-Economic Panel genetic subsample (SOEP-G).** The data collection for this study received ethical approval by the Vrije Universiteit Amsterdam, School of Business and Economics (Application No. 20181018.1.pkr730) and by the Max Planck Society (Application No. 2019_16).

**Berlin Aging Study II (BASE-II).** All participants gave written informed consent. All assessments were conducted in accordance with the Declaration of Helsinki and approved by the Ethics Committee of the Charité — Universitätsmedizin Berlin (approval numbers EA2/029/09 and EA2/144/16). BASE-II is registered in the German Clinical Trials Registry as DRKS00009277.

## Acknowledgements

The SHIP authors are grateful to Holger Prokisch and Thomas Meitinger (Helmholtz Zentrum München) for the genotyping of the SHIP-Trend cohort. The TwinLife authors thank Shirin Zare for performing the DNA extraction and Friederike David for sharing code for the PGS calculation. We thank the participants of SHIP, TwinLife, the Socio-Economic Panel, and the Berlin Aging Study II, without whom this work would not have been possible.

## Author Contributions

DF and LR conceived the study and designed the analyses. DF curated the genetic and phenotypic data and constructed the polygenic indices. DF performed the statistical analyses. AM performed statistical replication of analyses. DF and LR wrote the original draft of the manuscript. All authors interpreted the results, revised the manuscript and approved the final version. LR supervised the work.

