## Supplemental material for "Genomics of Educational Attainment Across 80 Years of Social and Political Transformation in Germany"

[1]Max Planck Research Group Biosocial — Biology, Social Disparities, and Development, Max Planck Institute for Human Development
[2]Department of Economics, School of Business and Economics, Vrije Universiteit Amsterdam
[3]DeSci Labs
[4]Max Planck Institute for Human Development
[5]SHARE Berlin Institute
[6]German Socio-Economic Panel Study Department
[7]Department of Psychology, Freie Universität Berlin
[8]Department of Social Sciences, Humboldt University
[9]Department of Psychology, Bielefeld University
[10]JICE — Joint Institute for Individualisation in a Changing Environment, Bielefeld University and University of Münster
[11]Institute of Human Genetics, University of Bonn, School of Medicine & University Hospital Bonn
[12]Institute of Neuroscience and Medicine (INM-1), Research Center Jülich
[13]Faculty of Sociology, Bielefeld University
[14]Department of Psychiatry and Psychotherapy, University Medicine Greifswald
[15]Institute for Community Medicine, University Medicine Greifswald
[16]Institute of Health Services Research in Dentistry, University of Münster
[17]Interfaculty Institute for Genetics and Functional Genomics, University Medicine Greifswald
[18]Lübeck Interdisciplinary Platform for Genome Analytics (LIGA), University of Lübeck
[19]Center for Lifespan Psychology, Max Planck Institute for Human Development
[20]Max Planck UCL Centre for Computational Psychiatry and Ageing Research
[21]Department of Psychology, Humboldt-Universität zu Berlin
[22]Institute of Gender in Medicine, Charité — Universitätsmedizin Berlin
[23]Department of Endocrinology and Metabolic Diseases, Charité — University Medical Center Berlin, corporate member of Freie Universität Berlin and Humboldt-Universität zu Berlin
[24]Berlin Institute of Health Center for Regenerative Therapies (BCRT), Charité — Universitätsmedizin Berlin, corporate member of Freie Universität Berlin, Humboldt-Universität zu Berlin, and Berlin Institute of Health
[25]Genetic Epidemiology, Department of Psychiatry, Amsterdam University Medical Centers
[26]Department of Economics, University of Bologna
[27]WZB Berlin Social Science Center
[28]Chair of Macroeconomics and Development, Goethe University Frankfurt
[29]Department of Psychology, The University of Texas at Austin
[30]Population Research Center, The University of Texas at Austin

[31]Department of Psychiatry, Amsterdam UMC, University of Amsterdam
[32]School of Biological and Behavioural Sciences, Queen Mary University of London

*Keywords*: Education, German Reunification, Educational Disparities, Polygenic Index, Gene–environment interactions

### Supplementary Methods

#### SNP-based heritability (GREML): implementation detail

Genetic relatedness matrices (GRMs) were built from harmonized autosomal SNPs (MAF ≥ .01) among approximately unrelated individuals, and restricted maximum likelihood (REML) partitioned phenotypic variance into an additive-SNP component and a residual to yield $h^2_{\text{SNP}}$. The primary stratification was the four-cell Region (East/West) × Time (pre/post-1990) design; the finer six-cell split was not fitted because two cells fell below the sample floor for stable REML.

Several implementation choices departed from the plan. HWE filtering was not applied at GRM construction, because GCTA's GRM routine exposes no HWE flag and pooled-sample HWE is confounded with cross-study allele-frequency structure. Nothing substituted for it at the variant level beyond the INFO ≥ 0.9 threshold, the cross-study SNP intersection, and the MAF ≥ .01 filter applied at GRM construction itself; pre-imputation quality control was performed by the contributing data providers and differed across studies. What a pooled HWE screen would guard against here is genotypes from three array families entering one relatedness matrix, and that is held apart by the construction of the primary estimator rather than by a variant filter, because its GRMs are built within study on within-study allele frequencies and combined block-diagonally under a categorical contributing-study fixed effect. Genotyping error arising within a study is not removed by that construction. Birth year (linear and quadratic, centred) entered the quantitative covariates alongside up to 20 ancestry PCs calculated across contributing studies, and gender entered the categorical covariates, so that broad secular birth-year trends in education were absorbed before variance partitioning. The primary cell-level estimator used block-diagonal GRMs assembled from study-specific GRMs (within-study allele frequencies, --grm-cutoff 0.05) with a categorical contributing-study fixed effect, so that cross-study structure and study mean differences did not load onto the genetic variance component; a pooled cross-study GRM at --grm-cutoff 0.20 was retained as a liberal-kinship sensitivity, relaxing the plan's 0.025 cutoff and single-GRM design. Statistical power was evaluated with a scripted implementation of the Visscher standard-error formula rather than the GCTA web calculator, which was unreachable from the compute environment (Visscher et al., 2014).

Heterogeneity in $h^2_{\text{SNP}}$ across the four cells ($H_0$: all cell heritabilities equal) was tested by an omnibus Cochran's $Q$ on the inverse-variance-weighted cell estimates, with pairwise $Z$-tests and an $I^2$ statistic, in place of the plan's joint multi-GRM LRT: GCTA's multi-GRM REML requires overlapping individuals across GRMs and does not support the disjoint-sample cell structure, whereas Cochran's $Q$ on the heritability ratios is the standard meta-analytic analogue and a closer match to the plan's ratio-equality $H_0$. The implied additive genetic variance $V_G = h^2_{\text{SNP}} \times V_P$ is reported per cell descriptively; a formal $Q$ test on $V_G$ was not run because the two imprecise post-1990 cells would dominate it. As exploratory complements, the same block-diagonal GREML was fitted for simpler groupings of the same sample — Region (East vs. West), recorded gender (male vs. female), and Region × gender — each with its own Cochran's $Q$; these surface group-level heterogeneity patterns rather than calibrated per-group population estimates, and recorded gender/sex labels index neither gender identity nor biological sex.

### Supplementary Results

#### Single-study (SOEP-G) test of the East–West educational attainment pattern

The East–West difference in the association between PGI-Education and educational attainment first reported in SOEP-G (Fraemke et al., 2025) was estimated with an ancestry adjustment specific to that study (residualizing the PGI on within-SOEP principal components), whereas the pooled multi-study pipeline here residualizes on shared ancestry principal components calculated across studies. To compare like with like, we restricted the current data to SOEP-G and reconstructed the earlier within-SOEP ancestry adjustment. The East German PGI-Education slope strengthened over birth year, from .10 (95% CI [−.09, .29]) among the earliest birth years to .50 (95% CI [.32, .69]) among the most recent, whereas the West German slope changed little, from .22 (95% CI [.12, .32]) to .35 (95% CI [.25, .45]; Figure S1). The per-region birth-year amplification was $b = .11$, 95% CI [.03, .19], $p = .007$ in East Germany and $b = .03$, 95% CI [−.01, .08], $p = .166$ in West Germany, and the PGI-Education × birth-year × region interaction was $b = .10$, 95% CI [.00, .19], $p = .048$ — a borderline East–West difference in the same direction as the earlier report (Fraemke et al., 2025).

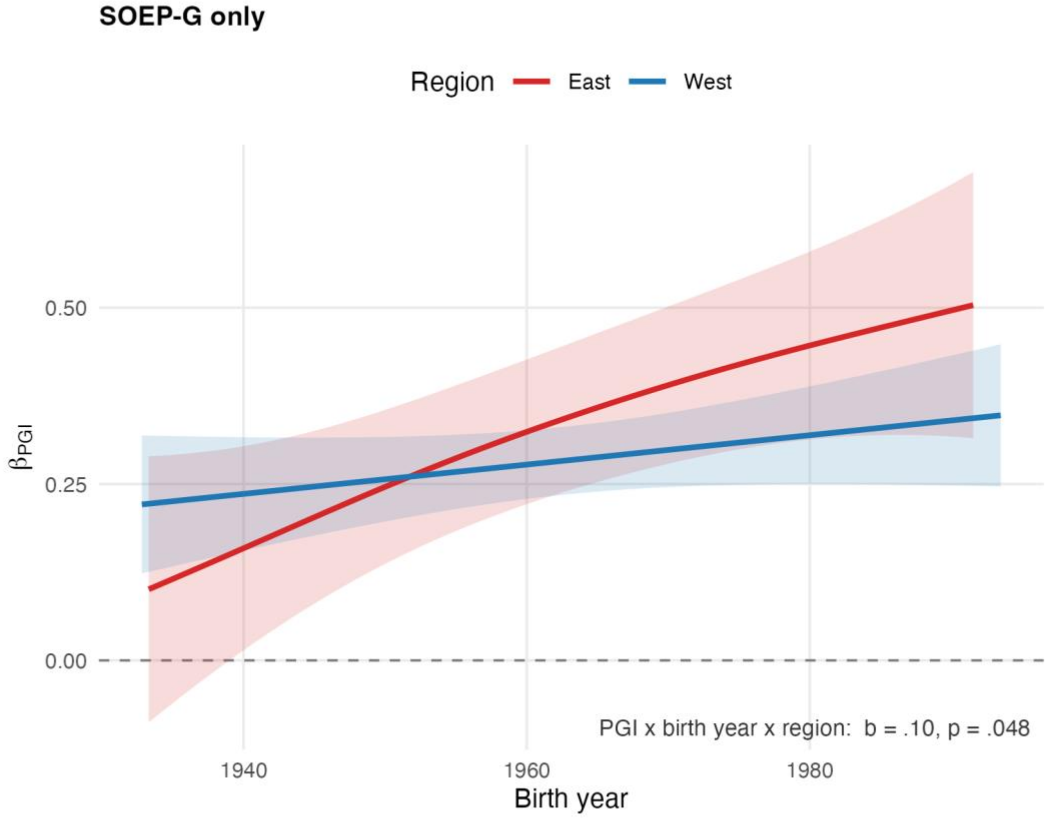

**Figure S1**

*Single-study (SOEP-G) association between PGI-Education and educational attainment over birth year, by region.*

Estimated within SOEP-G only, using the within-SOEP ancestry adjustment of the earlier single-study report (the PGI residualized on the within-SOEP principal components rather than the pooled pipeline's across-study components). Red lines are East Germany and blue lines are West Germany, the colour convention used throughout this article; they are the model-based PGI-

Education slopes for relative years of education over birth year, with 95% confidence-interval ribbons. The PGI-Education × birth-year × region interaction is reported in the text. The model is gender-pooled. Slopes are per 1 *SD* of PGI-Education. SOEP-G analytic sample, $n$ = 1,981 (East $n$ = 444; West $n$ = 1,537).

**Migration-composition diagnostic**

The planned migration-composition analysis was not interpreted as a sensitivity analysis of the present pooled multi-study findings. It was designed to examine the previously reported SOEP-G East–West difference in the association between PGI-Education and educational attainment (Fraemke et al., 2025); the corresponding regional moderation was not statistically distinguishable from zero in the combined four-study analyses, so migration composition was not pursued as an explanation of the current findings (Supplementary Data S3). Restricting the diagnostic to SOEP-G alone would leave the prespecified test substantially underpowered. The question has in any case already been examined in that sample, where PGI-Education did not differ between East and West Germany either before or after reunification, and where the report itself noted that firmer effect-size estimates would require larger German genetic datasets (Fraemke et al., 2025).

**Leave-one-study-out robustness**

To assess whether any single study drives the focal regional, birth-year, and gender associations, we re-estimated each focal model while omitting one contributing study at a time (leave-one-study-out). The educational attainment folds omit SOEP, TwinLife, BASE-II, or SHIP; the educational mobility folds omit SOEP, TwinLife, or BASE-II (SHIP does not contribute to the educational mobility subsample). Figure S2, Figure S3, and Figure S4 reproduce the focal educational attainment, educational mobility, and gender figures under each omission, with one panel per omitted study. In every panel the coloured curve or point is the leave-one-out estimate with its 95% confidence band; comparing panels shows whether the associations are stable across study omissions. Table S1 reports the composition of each reduced sample.

The associations were broadly stable across omissions, with two qualifications, set out below. For educational attainment, the association between PGI-Education and educational attainment stayed positive across the supported birth-year range in every omission, in both regions and both genders, and the linear birth-year specification held throughout, with no reduced sample showing evidence of nonlinearity (all leave-one-out region tests were non-significant, the strongest reaching only $p$ = .280; Figure S2, Figure S4, blocks A and C).

For educational mobility, the overall association remained positive in both regions under every omission, and East Germany exceeded West Germany in all cases (Figure S3, panel A). The East–West gap was widest when SOEP was omitted (East slope = .22, 95% CI [.17, .27]; West slope = .10, 95% CI [.06, .13]) and narrowest when BASE-II was omitted (East slope = .20, 95% CI [.14, .26]; West slope = .17, 95% CI [.14, .20]). The nonlinear birth-year specification that was marginally supported in the full educational mobility sample ($p$ = .041) was not retained in any reduced sample (all leave-one-out tests were non-significant, the strongest residual signal $p$ = .231; Figure S3, panel B; Figure S4, blocks B and D). The male–female difference in the educational mobility slope excluded zero only within the narrow, study-dependent birth-year windows shaded in Figure S4, block D.

Because omitting a study also lowers statistical power and narrows birth-year coverage (Table S1), these fluctuations in magnitude cannot be attributed to any single study

(Supplementary Data S4 and S7). Per-study estimates are given in the 01_PerStudy_region sheets of Supplementary Data S4 and S7, which fit the region-varying model of the PGI-Education association over birth year within each contributing study separately. Those fits are diagnostic rather than a parallel analysis. Each study covers only part of the birth-year range, so a per-study curve is estimated over a fraction of the span the combined analysis covers, and the studies differ in regional composition.

The East–West difference in the association between parental education and educational attainment, reported in the main text, was re-estimated over the same educational mobility folds. It was negative and statistically significant in every fold, but its magnitude varied. Against a full-sample contrast of $b = -0.216$, 95% CI [−0.278, −0.154], omitting BASE-II reduced it to $b = -0.072$, 95% CI [−0.138, −0.006], $p = .032$, whereas omitting SOEP widened it to $b = -0.303$, 95% CI [−0.372, −0.234] and omitting TwinLife gave $b = -0.131$, 95% CI [−0.216, −0.047]. The per-fold regional estimates locate the sensitivity. The East German association rises from .39 to .58 when BASE-II is omitted, so BASE-II's East German participants carry much of the difference. BASE-II recruited in and around Berlin, a mainly urban area, through a non-random scheme that over-represents participants with higher education and higher life satisfaction (Bertram et al., 2014), so its East German participants are not a general sample of East Germany. The regional difference therefore held in direction across omissions, while its size depended on which studies contributed (Supplementary Data S6).

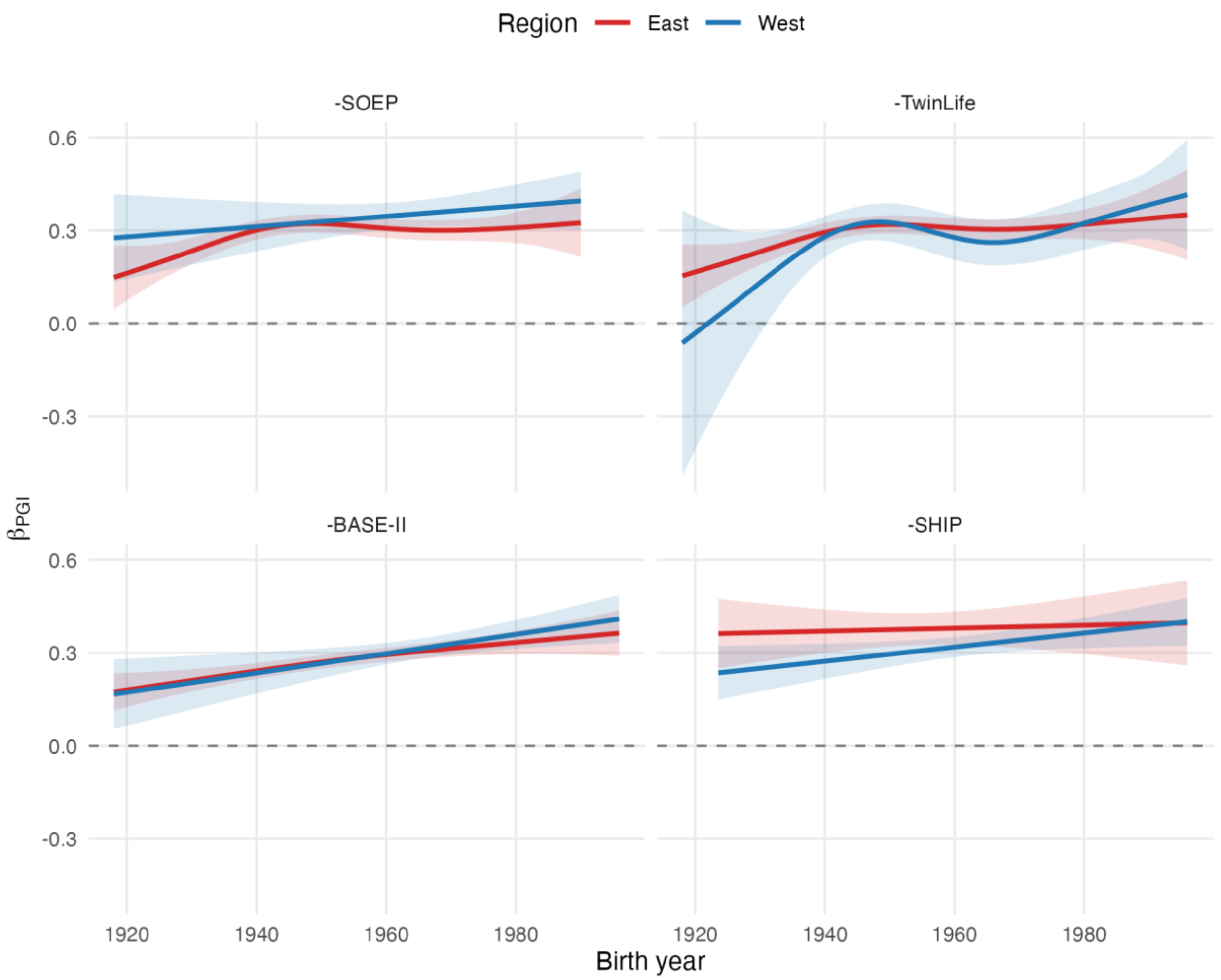

**Figure S2**

*Leave-one-study-out robustness of the association between PGI-Education and educational attainment, by region.*

Each panel omits one contributing study (labelled). Red lines are East Germany and blue lines are West Germany, the colour convention used throughout this article; they are the model-based PGI-Education slopes for educational attainment over birth year estimated with that study left out, with 95% confidence-interval ribbons. The gender-pooled region model is shown. Slopes are per 1 *SD* of PGI-Education. Sample size differs by fold; the per-fold composition is given in Supplementary Table S1.

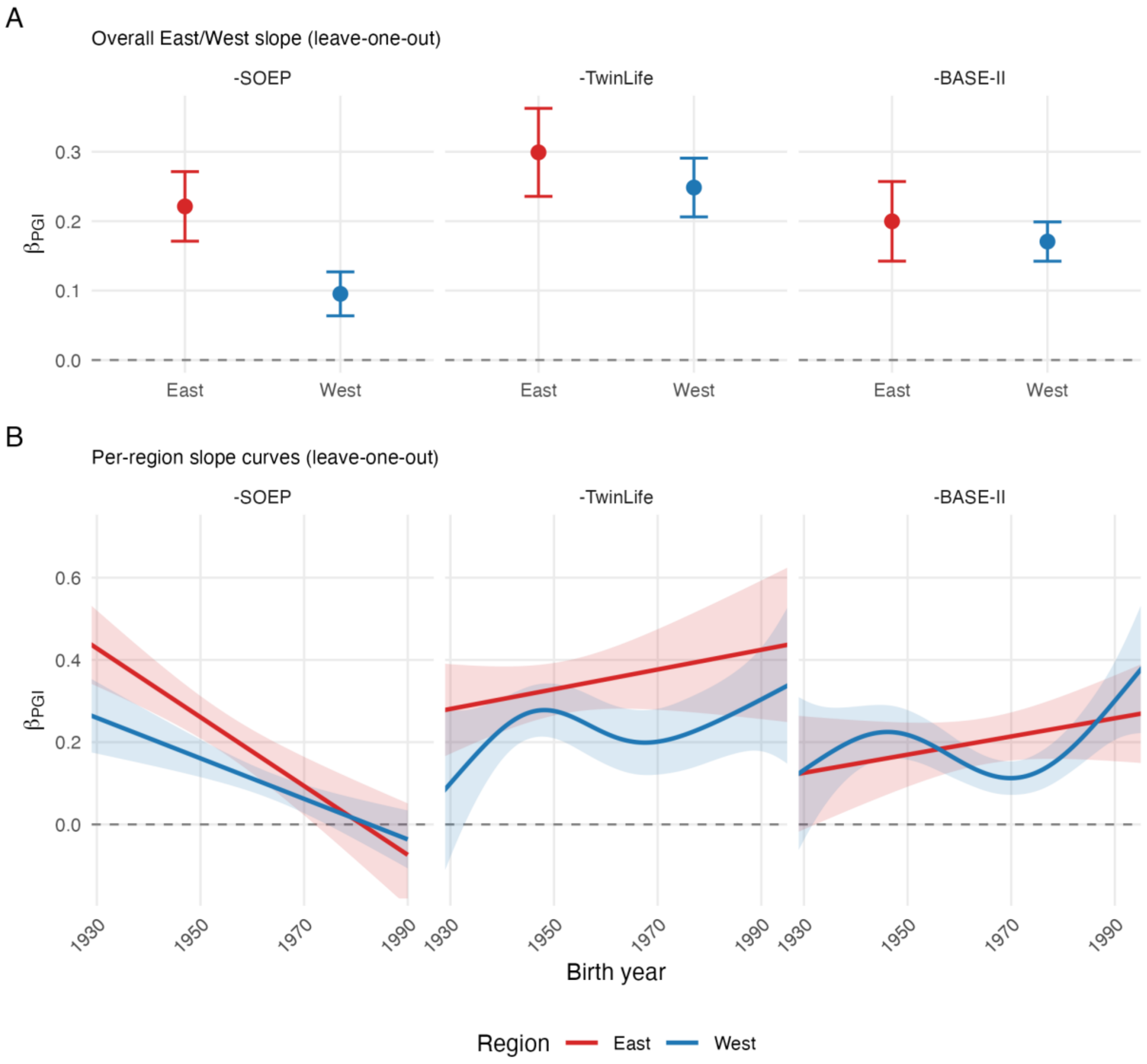

**Figure S3**

*Leave-one-study-out robustness of the association between PGI-Education and educational mobility, by region.*

Each panel omits one contributing study (labelled). Panel A shows the overall East and West PGI-Education slopes for educational mobility as points with 95% confidence-interval whiskers. Panel B shows the per-region slope curves over birth year, as leave-one-out estimates with 95% confidence-interval ribbons. Red denotes East Germany and blue denotes West Germany in both panels, the colour convention used throughout this article. Only the region curves are shown; the East-minus-West difference-smooth band from the corresponding main-text figure has no per-fold frozen export and is omitted here. Slopes are per 1 *SD* of PGI-Education. Sample size differs by fold; the per-fold composition is given in Supplementary Table S1.

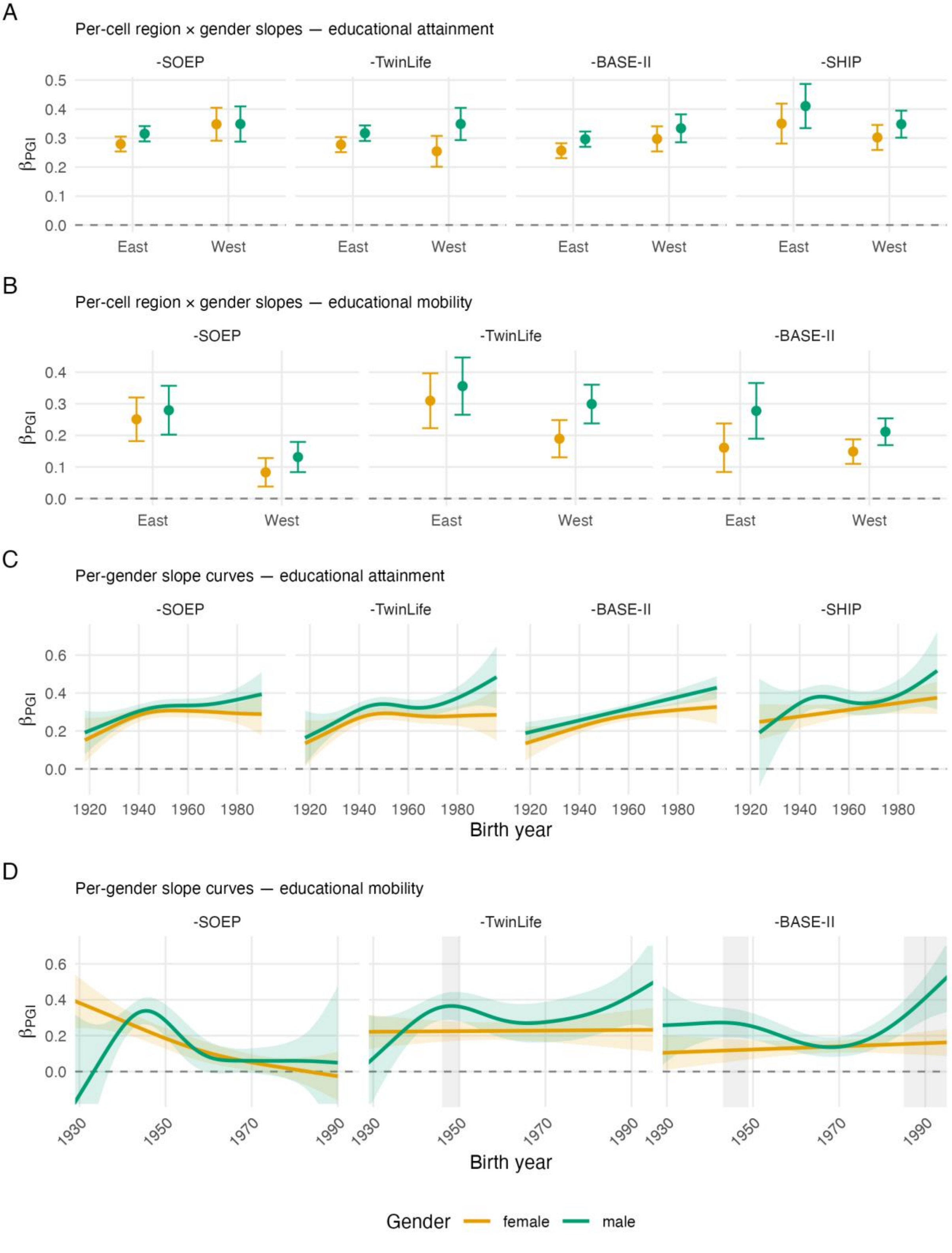
A
Per-cell region × gender slopes — educational attainment
-SOEP
-TwinLife
-BASE-II
-SHIP
βPGI
East
West
B
Per-cell region × gender slopes — educational mobility
C
Per-gender slope curves — educational attainment
Birth year
D
Per-gender slope curves — educational mobility
Gender
female
male

**Figure S4**

*Leave-one-study-out robustness of gender differences in associations between PGI-Education and educational attainment and educational mobility.*

Leave-one-study-out analogue of Figure 4 in the main text; each panel omits one contributing study (labelled). Blocks A and B show the per-cell region × gender PGI-Education slopes (points with 95% confidence-interval whiskers, dodged within region) for educational attainment and educational mobility. Blocks C and D show the region-pooled per-gender PGI-Education slope curves over birth year (one line per gender, with 95% confidence-interval ribbons) for educational attainment and educational mobility; in block D the grey vertical bands mark the birth-year windows where the male–female difference's 95% simultaneous confidence interval excludes zero. Amber denotes women and green denotes men throughout, the colour convention used in the main text. Slopes are per 1 *SD* of PGI-Education. Sample size differs by fold; the per-fold composition is given in Supplementary Table S1.

**Table S1**

*Sample composition is shown under each leave-one-study-out omission.*

| Sample | N | Mean birth year | Mean years of education | Mean PGI-Education (z) | % East | % female |
|---|---|---|---|---|---|---|
| Full sample | 13,049 | 1956.2 | 12.97 | 0.00 | 70.9 | 52.7 |
| -SOEP | 11,068 | 1955.3 | 12.95 | −0.01 | 79.6 | 52.5 |
| -TwinLife | 11,034 | 1953.6 | 12.69 | 0.00 | 81.0 | 51.7 |
| -BASE-II | 11,919 | 1957.5 | 12.85 | −0.04 | 72.9 | 53.0 |
| -SHIP | 5,126 | 1960.8 | 13.87 | 0.11 | 26.0 | 55.1 |

*Note.* Each row omits the labelled study from the combined sample; "Full sample" retains all contributing studies. *N* is the number of genotyped participants contributing to the educational attainment analyses. PGI-Education is standardized in the full sample. Educational-mobility analyses use a narrower subsample and omit SHIP.

### Height negative control

A negative control asks whether a design reproduces the pattern of interest when the predictor has no substantive reason to produce it (Lipsitch et al., 2010). PGI-Height was constructed, residualized on ancestry principal components, and standardized exactly as PGI-Education was, so it carries the same exposure to population stratification and to cohort differences in genotyping, but there is no substantive reason for its association with the educational outcomes to strengthen over birth year or to differ by region or gender. Those moderation terms are what the control was prespecified to test.

The prespecified height negative-control analyses examined whether linear counterparts of the focal birth-year, regional, and gender patterns appeared when PGI-Education was replaced by PGI-Height. None did. The association with educational attainment showed no evidence of increasing over birth year ($b = -.006$, 95% CI [−.028, .016], $p = .581$), there was no East–West difference in intergenerational educational mobility ($b = .021$, 95% CI [−.053, .095], $p = .581$), and neither gender term was distinguishable from zero (educational attainment $b = .030$, 95% CI [−.013, .073], $p = .166$; educational mobility $b = -.001$, 95% CI [−.074, .073], $p = .986$).

As expected, PGI-Height was strongly associated with height ($b$ = .412, 95% CI [.399, .424], $p$ < .001). That association showed no evidence of changing over birth year ($b$ = .010, 95% CI [−.003, .022], $p$ = .130), of differing by region, or of differing by gender (Supplementary Data S3 and S6). The height negative control therefore did not reproduce the focal linear patterns in the educational outcomes.

**SNP-based heritability by region and era**

The main text reports SNP-based heritability of educational attainment estimated by GREML within Region × Time cells, by region, and by gender. Figure S5 displays those estimates with their confidence intervals and heterogeneity tests. Implementation detail is given in the Supplementary Methods.

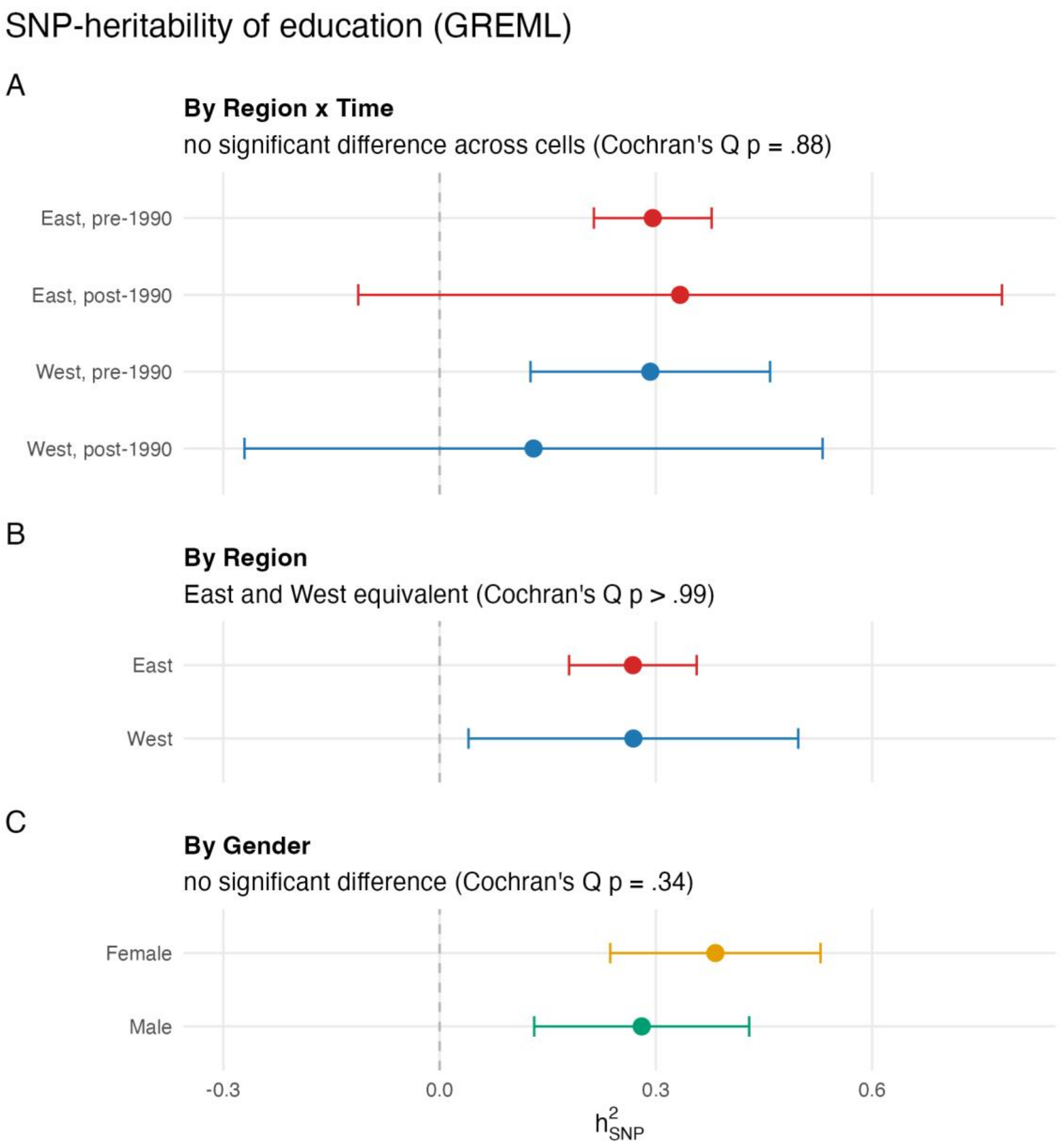

**Figure S5**
*SNP-based heritability of educational attainment by region and era.*

GREML estimates on the unrelated-individual subset. Panel A shows the four Region × Time cells, panel B the two regions, and panel C the two gender groups, each with 95% confidence intervals and the corresponding Cochran's $Q$ verdict annotated. Red denotes East Germany and blue denotes West Germany; amber denotes women and green denotes men, the colour conventions used throughout this article. The post-1990 cells in panel A are estimated on substantially smaller subsamples (East $n$ = 1,170, West $n$ = 1,423, against East $n$ = 7,111 and West $n$ = 2,993 before 1990) and their intervals are correspondingly wide; the absence of a detected difference should not be read as evidence that the heritabilities are equal. The unrelated-individual subset totals $n$ = 12,697 across cells.

**Gender difference across the PGI distribution**

Gender differences in educational attainment were evident in its mean, its variance, and its association with PGI-Education. The mean difference varied by birth-year group and reversed over time — men attained more than women among the oldest participants, converging in the middle birth-year group and reversing to a female advantage among the youngest (main-text descriptive results). The variance of educational attainment was consistently higher among men, with a male-to-female variance ratio of 1.10 on the years scale and 1.18 on the birth-year-standardized scale (both Fligner–Killeen $p < .001$), and men's variance exceeded women's within every birth-year group. Against this backdrop, PGI-Education was more strongly associated with educational attainment among men than women, as reported in the main text. Because differences in outcome dispersion can affect comparisons of association strength, we examined whether the gender difference persisted under dispersion-aware and scale-free specifications.

In an additional exploratory analysis, we examined whether the gender difference in the association between PGI-Education and educational attainment was concentrated at lower or higher values of PGI-Education. A single linear PGI-Education × gender interaction constrains the gap to change at a constant rate across PGI and cannot localize it, so we split the educational attainment sample into PGI-Education quintiles (pooled across region and contributing study) and compared the raw male and female means of relative years of education within each quintile (main-text Figure 4, panel B; Figure S6). At the lowest quintile, the male–female difference was small (0.05 *SD*, *SE* 0.03). The difference was larger in the two highest quintiles, where it was 0.18 and 0.15 *SD*; no quintile showed a female advantage. The female mean continued to rise into the highest quintile (0.38 *SD*), so higher-PGI women also attained relatively high educational positions. Descriptively, the gender difference in the association between PGI-Education and educational attainment was concentrated among higher-PGI individuals. Whether this gap changed across birth cohorts was not examined at the quintile level. The model-based counterpart is the PGI-Education × birth year × gender interaction, which is reported in the main text for both educational outcomes.

Whether the weaker female association merely reflects women's lower variance in educational attainment is addressed directly by the location-scale sensitivity analyses reported in the main text, in which the pooled PGI-Education × gender interactions remained distinguishable from zero when residual dispersion was modelled. Two assumption-light checks concur. The Spearman rank correlation — invariant to monotonic rescaling and less sensitive to the bounded, coarse education scale — was higher among men ($\rho = .31$, 95% CI [.29, .33]) than among women ($\rho = .27$, 95% CI [.25, .29]; Fisher-$z$ test $p = .023$), and an ordered-probit PGI-Education × gender interaction adjusted for birth year and region, which treated raw education as ordered categories rather than a continuous outcome, pointed the same way ($b = 0.046$, $p = .011$). Taken together, the weaker association among women was not eliminated by modelling residual dispersion or by using rank- and ordinal-scale checks. The latter checks were exploratory and did not include the family random intercept used in the prespecified models.

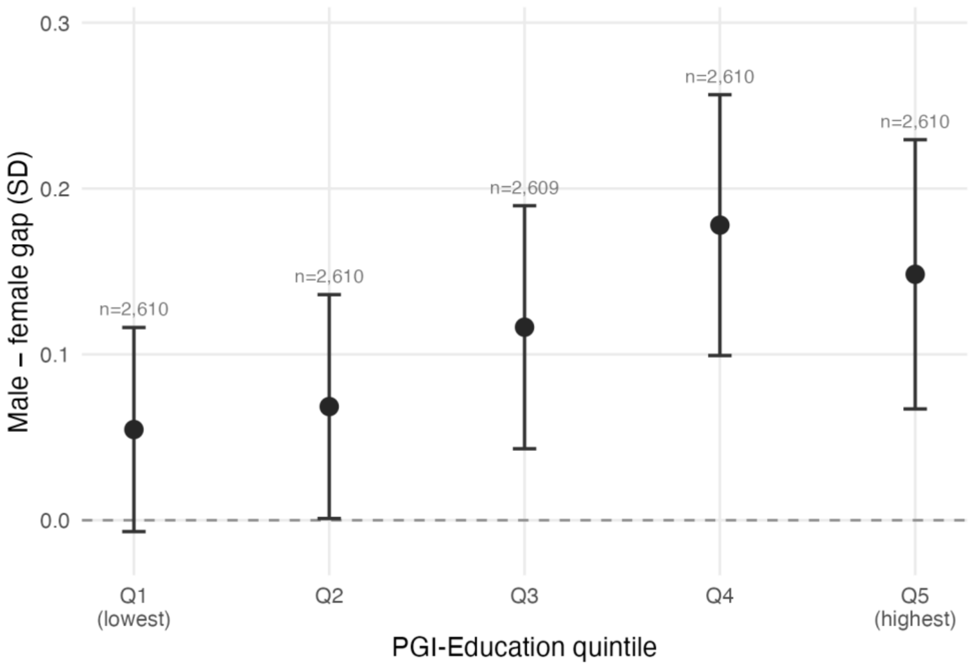

**Figure S6**

*Male–female gap in educational attainment across the PGI-Education distribution.*

Exploratory, descriptive; no model is fitted. The educational attainment sample is split into PGI-Education quintiles, pooled across region and contributing study (quintile 1 = lowest PGI). Points are the male-minus-female difference in mean relative years of education within each quintile, with 95% confidence intervals; *n* is the number of participants per quintile. The dashed horizontal line marks no gender difference. Educational attainment is in *SD* units. The per-gender means behind these differences are shown in main-text Figure 4, panel B. Educational attainment *n* = 13,049 (women *n* = 6,882; men *n* = 6,167).

**Parental education as a moderator of polygenic prediction**

Morris et al. (2026) reported that the association between PGI-Education and educational attainment in three British birth cohorts was disproportionately stronger among participants from more socioeconomically advantaged origins. Because the educational mobility models already carry birth-year-standardized parental education and its interactions with PGI-Education, region, and birth year, an equivalent test is available here. This analysis was not prespecified and is exploratory.

Because educational mobility is the difference between the respondent's and the parental birth-year-standardized score while parental education is itself a model term, the estimates below are numerically identical for educational attainment and for educational mobility; they are not specific to educational mobility.

Pooled across regions, parental education did not moderate the association between PGI-Education and educational outcomes ($b$ = 0.010, 95% CI [−0.020, 0.040], $p$ = .528). The moderation did, however, differ between East and West Germany ($b$ = 0.068, 95% CI [0.008, 0.127], $p$ = .025). Descriptively, the PGI-Education slope rose across the parental-education distribution in East Germany, from 0.22 at one standard deviation below the mean to 0.31 at one standard deviation above it, and declined in West Germany over the same range, from 0.23 to 0.18 (Figure S7). Neither region's moderation slope was individually distinguishable from zero

(East $b = 0.044$, 95% CI [−0.009, 0.096], $p = .104$; West $b = -0.024$, 95% CI [−0.053, 0.004], $p = .097$), so the regional contrast rests on the difference between two slopes that were not separately detected.

Parental education itself predicted educational attainment more strongly in West than in East Germany ($b = -0.218$, 95% CI [−0.281, −0.155], $p < .001$ for the East–West difference). In Figure S7, panel A, this gradient is what sets how far apart the lines lie within each region; the moderation under test is the change in that spacing across the PGI-Education axis.

The regional difference should be interpreted with care. Its direction is the opposite of a resource-amplification account, since West Germany — which retained early selection into stratified tracks and most closely resembles the British setting — showed no amplification, whereas the positive slope appeared in East Germany, where higher parental education was for part of the period a disadvantage in admission to higher education because access was preferentially extended to students from working-class and agricultural families and was additionally regulated by political criteria (Below, 2017; Betthäuser, 2019; Klein et al., 2019). Parental education in East Germany therefore does not index socioeconomic advantage in the sense intended in the British analyses. Given that neither regional slope was individually detected, and that this sample is substantially smaller than the British cohorts, we do not draw a substantive conclusion from the contrast.

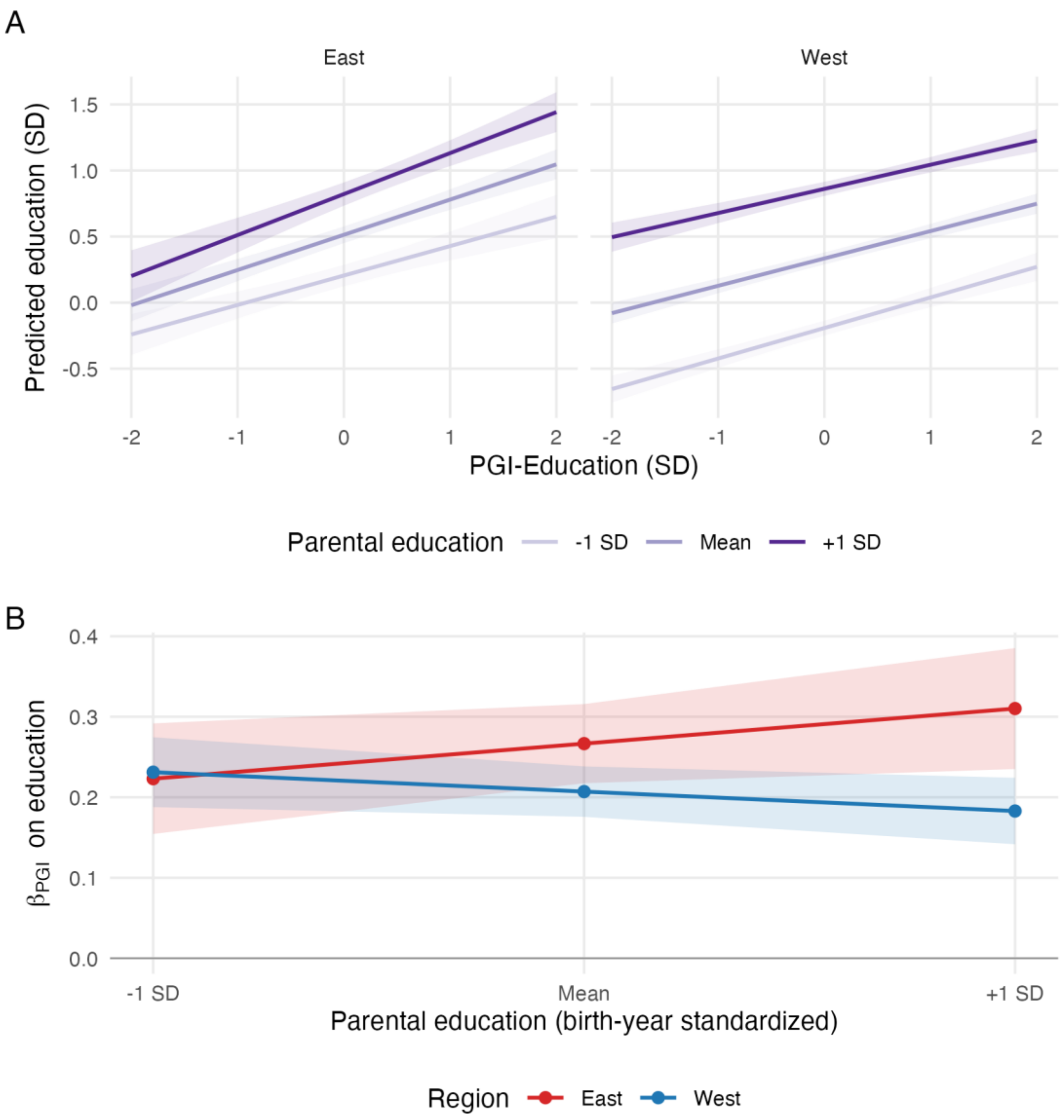

**Figure S7**

*Parental education as a moderator of PGI-Education prediction, by region.*

Estimated in the educational mobility analytic sample, which requires parental education. Panel A shows model-implied predicted educational attainment as a function of PGI-Education at one standard deviation below the mean, at the mean, and one standard deviation above the mean of birth-year-standardized parental education, separately for East and West Germany; non-parallel lines within a panel represent the interaction. Panel B shows the same interaction expressed as the PGI-Education slope across the parental-education distribution. In panel A the two regions are separate facets, and the three lines within each facet are shaded in a light-to-dark purple ramp running from parental education one standard deviation below the mean, through the mean, to one standard deviation above it. In panel B red denotes East Germany and blue denotes West Germany, the colour convention used throughout this article. Bands are 95% confidence intervals and slopes are per 1 *SD* of PGI-Education. Educational mobility analytic sample, $n$ = 4,904.

Panel A plots fitted values from the full model, evaluated at the sample mean of birth year and gender; because both are centred, the plotted lines represent the whole analytic sample rather than a subgroup, and are not a bivariate regression of educational attainment on PGI-Education. The model carries a single linear PGI-Education by parental-education term per region, so the lines are linear and evenly spaced by construction and are not independent estimates of a shape. How far apart the lines lie within a panel reflects the association of parental education with educational attainment, which is stronger in West Germany; the moderation under test is the change in that spacing across the PGI-Education axis. Because educational mobility is the difference between respondent and parental birth-year-standardized scores and parental education is a model term, these estimates are identical for educational attainment and educational mobility. The analysis is exploratory and was not prespecified.

**Outcome scale and estimator**

Relative years of education is bounded at both ends and coarsely graded, so an association estimated on it could in principle reflect the scale rather than the outcome. We re-estimated two focal terms, the PGI-Education × birth year interaction in educational attainment and the PGI-Education × region interaction in intergenerational educational mobility, changing one feature of the published model at a time — the outcome in raw years of education; the outcome replaced by its rank-based inverse normal transform, with the published covariates unchanged; and an ordered probit on the respondent's education categories, with the published right-hand side. The ordered probit cannot carry the per-family random intercept the other specifications use, so it was fitted without one and with standard errors clustered by family. These analyses were not part of the prespecified analysis plan.

The strengthening of the PGI-Education association with educational attainment over birth year held under all three changes (Table S2). On raw years of education the interaction was $b = 0.073$ years per standard deviation of PGI-Education, 95% CI [0.022, 0.124], $p = .005$, and its size relative to the PGI-Education main effect of the same model was 0.087 against 0.094 in the published model, so the birth-year standardization of the outcome moves it little.

The East–West difference in the PGI-Education association with intergenerational educational mobility kept its sign under all three changes and remained distinguishable from zero on raw years of education and under the rank transform. Under the ordered probit it was not ($b = 0.060$, 95% CI [−0.016, 0.137], cluster-robust $p = .121$; model-based $p = .093$). Of the three changes, only the change of estimator removed the contrast. The ordered probit enters the step from 13 to 18 years of education as one ordered category rather than as five years, so the tertiary transition no longer carries the weight its duration gives it on the years scale, and PGI-Education is particularly sensitive to that transition because the discovery signal it is built on is largely a college-completion contrast (Okbay et al., 2022). The same ordered probit detected the PGI-Education × gender interaction reported above ($p = .011$), so the estimator is not simply insensitive to interactions of this size in these data.

Differential censoring of the education measure does not account for the regional difference either. Within the educational mobility subsample, East German participants were at the ceiling of the measure more often than West German participants (27.3% against 24.4%) and at the floor less often (1.6% against 2.3%). Under the usual mechanism, by which censoring attenuates an association in the more censored group, that asymmetry would understate rather than produce the East–West difference. These are descriptive shares and no test was applied to them.

**Table S2**

*Outcome scale and estimator sensitivity are shown for the two focal interactions.*

| Contrast | Specification | *b* [95% CI] | *p* | Ratio to PGI-Education main effect |
|---|---|---|---|---|
| Educational attainment: PGI-Education × birth year | As published | .029 [.010, .048] | .002 | 0.094 |
| | Raw years of education | 0.073 [0.022, 0.124] | .005 | 0.087 |
| | Rank-based inverse normal transform | .023 [.004, .042] | .019 | 0.074 |
| | Ordered probit, no family random intercept | 0.032 [0.009, 0.056] | .006 | 0.093 |
| Intergenerational educational mobility: PGI-Education × region | As published | .073 [.017, .129] | .011 | 0.321 |
| | Raw years of education | 0.187 [0.036, 0.338] | .015 | 0.329 |
| | Rank-based inverse normal transform | .072 [.014, .130] | .014 | 0.314 |
| | Ordered probit, no family random intercept | 0.060 [−0.016, 0.137] | .121 | 0.214 |

*Note.* Each row changes one feature of the published model at the head of its block and is otherwise identical to it. Estimates are on different scales and are not comparable in size across rows — standard deviations of the birth-year-standardized outcome, years of education, normal scores, and latent-probit units — so the last column gives each interaction relative to the PGI-Education main effect of the same model, which is scale-free. The ordered probit cannot carry the per-family random intercept the other rows use; its *p* value comes from standard errors clustered by family, and the model-based value is given in the text. These analyses were not part of the prespecified analysis plan.

**Scope of the ancestry adjustment of PGI-Education**

The single-study analysis reported above differs from the pooled analysis in two ways at once, the sample and the scope of the ancestry adjustment, and the two can be separated. The published models residualize PGI-Education on ancestry principal components calculated across the contributing studies. We refitted the published model with PGI-Education residualized instead on principal components calculated within each contributing study, SHIP-START and SHIP-TREND separately because each carries its own principal-component analysis. Residualizing within a group also removes that group's mean. The specification was otherwise unchanged. Taken with the two single-study estimates, this completes a 2 × 2 of sample by scope of adjustment (Table S3). This analysis was not part of the prespecified analysis plan.

In the pooled four-study sample the PGI-Education × birth year × region interaction was not distinguishable from zero under either adjustment, $b = -.003$, $p = .879$ across studies and $b = -.008$, $p = .668$ within contributing study. Restricting the sample to SOEP-G moved the estimate much further, to $b = .081$, $p = .086$ under the across-study adjustment and $b = .097$, $p = .048$ under the within-study one. The divergence from the earlier single-study report therefore follows from the sample rather than from the scope of the ancestry adjustment.

Only the pooled pair changes one factor. The two single-study estimates carry no per-family random intercept and standardize PGI-Education within SOEP-G rather than over the pooled sample, so part of the difference between them is rescaling rather than adjustment, and the four estimates should not be read as one homogeneous grid.

**Table S3**

*The sample and scope of the ancestry adjustment of PGI-Education are shown as a 2 × 2.*

| Sample | Ancestry principal components | *n* | Family random intercept | *b* [95% CI] | *p* |
|---|---|---|---|---|---|
| Pooled, four studies | Across studies (published) | 13,049 | Yes | −.003 [−.040, .035] | .879 |
| Pooled, four studies | Within contributing study | 13,049 | Yes | −.008 [−.046, .029] | .668 |
| SOEP-G only | Across studies | 1,981 | No | .081 [−.011, .174] | .086 |
| SOEP-G only | Within SOEP-G | 1,981 | No | .097 [.001, .193] | .048 |

*Note.* Each row gives the PGI-Education × birth year × region interaction for educational attainment. The first row is the published specification and the second changes only the scope of the ancestry residualization, which is calculated within each contributing study, with SHIP-START and SHIP-TREND taken separately because each carries its own principal-component analysis, and which therefore also removes each group's mean. Pooling SHIP-START and SHIP-TREND into one group instead moves the estimate by 0.0002. The two single-study rows are read from the same frozen analysis as Supplementary Figure S1; they are not fit-identical to the pooled pair, because they carry no per-family random intercept and standardize PGI-Education within SOEP-G rather than over the pooled sample, so part of the difference between them is rescaling rather than adjustment. Only the pooled pair changes one factor. These analyses were not part of the prespecified analysis plan.

**Ancestry adjustment of the outcome model**

The published models residualize PGI-Education on ancestry principal components before fitting, which removes the components' linear association with the predictor but leaves the outcome model itself unadjusted for them. We refitted the four models the manuscript reports its headline interactions from, first adding the ten components as main effects in the outcome model, then adding their interactions with birth year, region, and gender. Each rung adds one block of terms to the rung above it; the main effects come first because an interaction with a principal component cannot be fitted without it. These analyses were not part of the prespecified analysis plan.

The adjustment was substantial rather than nominal (Table S4). In every model the ten principal-component main effects were jointly distinguishable from zero, and so were their interactions with birth year, region, and gender, all $p < .001$. Ancestry axes therefore predicted both educational outcomes directly and interacted with birth year, region, and gender in this sample.

The four focal interactions were nevertheless insensitive to that adjustment. No estimate changed sign, and every adjusted standard error was smaller than its published counterpart, so the additional terms did not cost precision in the focal contrasts — consistent with the component block explaining enough outcome variance to offset the degrees of freedom it consumes. Under the full adjustment the birth-year amplification in educational attainment and the gender difference in educational attainment were somewhat stronger than as published, while the regional contrast in educational mobility and the gender difference in educational mobility did not materially move.

**Table S4**

*The ancestry adjustment of the outcome model is shown for the four reported PGI-Education × moderator interactions.*

| Finding | Specification | *b* [95% CI] | *SE* (change) | *p* | Coefficients | Test of the block added |
|---|---|---|---|---|---|---|
| Educational attainment: PGI-Education × birth year | As published | .0292 [.0105, .0480] | 0.0096 | .002 | 15 | — |
| | + PC main effects | .0309 [.0121, .0496] | 0.0095 (−0.4%) | .001 | 25 | $\chi^2(10) = 150.3$, $p < .001$ |
| | + PC × moderators | .0328 [.0141, .0514] | 0.0095 (−0.7%) | < .001 | 55 | $\chi^2(30) = 203.0$, $p < .001$ |
| Intergenerational educational mobility: PGI-Education × region | As published | .0727 [.0166, .1289] | 0.0286 | .011 | 13 | — |
| | + PC main effects | .0668 [.0115, .1221] | 0.0282 (−1.4%) | .018 | 23 | $\chi^2(10) = 168.5$, $p < .001$ |
| | + PC × moderators | .0712 [.0158, .1266] | 0.0283 (−1.3%) | .012 | 53 | $\chi^2(30) = 99.8$, $p < .001$ |
| Educational attainment: PGI-Education × gender | As published | −.0388 [−.0762, −.0014] | 0.0191 | .042 | 16 | — |
| | + PC main effects | −.0373 [−.0746, −.0001] | 0.0190 (−0.5%) | .049 | 26 | $\chi^2(10) = 150.5$, $p < .001$ |
| | + PC × moderators | −.0410 [−.0781, −.0039] | 0.0189 (−1.0%) | .030 | 56 | $\chi^2(30) = 202.9$, $p < .001$ |
| Intergenerational educational mobility: PGI-Education × gender | As published | −.0656 [−.1238, −.0074] | 0.0297 | .027 | 24 | — |
| | + PC main effects | −.0643 [−.1217, −.0070] | 0.0293 (−1.5%) | .028 | 34 | $\chi^2(10) = 166.2$, $p < .001$ |
| | + PC × moderators | −.0638 [−.1210, −.0065] | 0.0292 (−1.6%) | .029 | 64 | $\chi^2(30) = 100.9$, $p < .001$ |

*Note.* Each finding is shown under the published model and under two further rungs, each adding one block of terms to the rung above it — the ten ancestry principal components as main effects in the outcome model, then their interactions with birth year, region, and gender. Main effects come first because an interaction with a principal component cannot be fitted without it. The standard error is followed by its change relative to the published model, and the last column is a joint Wald test of the block added at that rung, Wald rather than likelihood-ratio because the log-likelihoods of these models are not comparable. A falling standard error does not by itself move *p*. It falls furthest in the two gender blocks, where *p* changes least, because the estimate shifts over the same rungs. Coefficients is the number of fixed-effect coefficients in the model. The published models already residualize PGI-Education on the same components. Educational attainment $n = 13{,}049$; intergenerational educational mobility $n = 4{,}904$. These analyses were not part of the prespecified analysis plan.
